# Chronic cisplatin exposure does not affect epimutations in *C. elegans* but induces fluctuations in tRNA-derived small non-coding RNAs

**DOI:** 10.1101/2023.07.13.548810

**Authors:** Manon Fallet, Rachel Wilson, Peter Sarkies

## Abstract

The individual lifestyle and environment of an organism can influence its phenotype and potentially the phenotype of its offspring. The different genetic and non-genetic components of the inheritance system and their mutual interactions are key mechanisms to generate inherited phenotypic changes. Epigenetic changes can be transmitted between generations independently from changes in DNA sequence. In *C. elegans*, epigenetic differences, i.e., epimutations, mediated by small non-coding RNAs, particularly 22G-RNAs, as well as chromatin have been identified and their average persistence is 3 to 5 generations. In addition, previous research showed that some epimutations had a longer duration and concerned genes that were enriched for multiple components of xenobiotic response pathways. These results raise the possibility that environmental stresses might change the rate at which epimutations occur, with potential significance for adaptation. In this work, we explore this question by propagating *C. elegans* lines either in control conditions or in moderate or high doses of cisplatin, which introduces genotoxic stress by damaging DNA. Our results show that cisplatin has a limited effect on global small non-coding RNAs epimutations and epimutations in gene expression levels. However, cisplatin exposure leads to increased fluctuations in the levels of small non-coding RNAs derived from tRNA cleavage. We show that changes in tRNA-derived small RNAs may be associated with gene expression changes. Our work shows that epimutations are not substantially altered by cisplatin exposure but identifies transient changes in tRNA-derived small RNAs as a potential source of transcriptional variation induced by genotoxic stress.

## Introduction

Environmental factors can influence the phenotype of organisms by interfering with genetic and non-genetic factors^1, 2^. Among non-genetic changes that can be brought about by environmental responses, epigenetics has been defined as changes in gene expression that are mitotically and/or meiotically inheritable without altering the DNA sequence^3^. Epigenetic alterations known as epimutations, that are transmitted to subsequent generations can be caused by several molecular mechanisms including small non-coding RNAs, DNA methylation, histone modifications and prions^4–8^. Epimutations may be triggered by environmental stimuli but can also arise spontaneously in the absence of a specific stimulus. In general, the observed epigenetic modifications only remain for a few generations, but they can sometimes last much longer^9–15^.

The nematode worm *C. elegans* is an ideal organism to study the scope of epimutations in multigenerational studies thanks to its short generation time. In *C. elegans*, transgenerationally inherited transcriptional silencing is mediated by small RNAs (sRNAs). This silencing is initiated by double-stranded RNA (dsRNA) that will activate RNA-induced epigenetic silencing (RNAe), when the dsRNA is from endogenous origins, or the RNA interference (RNAi) pathway if the dsRNA is from exogenous origins^16^. Both RNA silencing systems have similar machinery: dsRNAs are processed into primary small-RNAs (sRNAs) with a length between 18 and 30 nucleotides. Argonaute proteins (AGOs) recognize the sRNAs and form RNA_induced silencing complexes (RISC). The targeting of transcripts by RISCs is followed by the cleavage of the targeted mRNA and the recruitment of an RNA-dependent RNA polymerase (RdRP) that will recognize the cleaved mRNA and generate secondary sRNAs. These secondary RNAs are antisense to the mRNA target, start with a guanine and have a length of 22 nucleotides (22G-RNAs). 22G-RNAs are loaded into a second class of AGOs, the Worm-specific Argonautes (WAGOs) that will repress the mRNA target expression at the transcriptional and post-transcriptional level^17^. 22G-RNAs and the associate AGO protein can be transmitted through generations via the gametes resulting in transgenerational persistence of the transcriptional silencing^18^. In addition to epigenetic memory through small non-coding RNAs transmission, epigenetic inheritance in *C. elegans* can also be mediated by other epigenetic factors such as changes in chromatin structure. Indeed, the chromatin is highly dynamic and from its organisation depend on gene expression regulation. For instance, in response to a thermal stress, a loss of the histone mark H3K9me3 was associated to a switch toward heterochromatin for specific genomic regions, leading to a long-lasting (14 generations) gene repression^19^.

Building on the detailed mechanistic understanding of transgenerational epigenetic inheritance in *C. elegans*, we have recently begun to address the contribution that these processes might make to evolution. Key to these experiments was the use of mutation accumulation lines, which are propagated under minimal selection to enable unbiased assessment of mutations and epimutations regardless of their effect on fitness^20^. These studies demonstrated that epimutations, defined as heritable changes in small non-coding RNAs, chromatin accessibility or gene expression occur in populations with a rate around 100 times greater than DNA sequence changes. However, epimutations were also ephemeral, lasting only around 4 generations on average^9, 11^.

Our previous experiments investigated epimutations in a stable environment. Classical evolutionary theory posits that DNA sequence changes are not affected in an adaptive sense by stress^21^. Whilst the overall rate or type of mutations is clearly affected by certain types of stimuli^22, 23^, there is little evidence that the types of genes affected by mutations changes such that mutations are more likely to occur in genes involved in stress responses. However, epimutations may behave differently. Our previous work demonstrated that epimutations are more likely to occur in genes involved in environmental responses, even in the absence of stress. Thus, defining how the properties of epimutations are affected by stress is key to understanding what role epimutations may play in evolutionary processes involving adaptation by natural selection.

In this study, we investigated the effect of genotoxic stress on epimutations. We constructed MA lines exposed to the DNA damaging agent cisplatin, which is already known to induce mutations, and investigated how the rate, spectrum, and stability of epimutations was affected. Our results indicated that continuous cisplatin exposure, despite inducing mutations in DNA sequence, had little effect on the occurrence or stability of epimutations, either at the level of gene expression or 22G-RNAs. However, cisplatin exposure led to induction of heritable changes in fragments of tRNA 3’ halves. Intriguingly, potential target genes showed expression changes associated with tRNA epimutations, suggesting that epimutations in tRNAs might have consequences for gene regulation. Overall, our work suggests that most types of epimutation behave according to the paradigm established for DNA sequence changes, but uncovers tRNA-derived small RNAs, previously implicated in epigenetic processes in mammals but not nematodes, as a new source of transcriptional variability within populations exposed to genotoxic stress.

## Methods

### Construction of mutation accumulation lines

Mutation Accumulation (MA) lines experiments in *C. elegans* have been described in previous literature^11, 24^. In brief, MA lines consist of minimizing the effective population size by selecting only the minimal number of individuals required to rise each following generation. According to a previous study^11^ two worms were selected to establish each new generation. The *Caenorhabditis elegans* (N2) worms were grown on Nematode Growth Medium (NGM) plates with an OP50 *Escherichia*. *coli* lawn and kept at 20 □C in a temperature-controlled incubator. Six independent lineages were constructed: two lineages in control conditions (lineages A (C1) and B (C2)), two lineages grown on plates enriched with 75μM of cisplatin (Low dose) (lineages C (L1) and D (L2)) and two lineages grown on plates enriched with 150μM of cisplatin (High dose) (lineages E (H1) and F (H2)). First, N2 worms were bleached to obtain a synchronized population (day 1). On day 2, L1 worms were seeded onto each of the 6 plates (Generation 0, Pre-accumulation line). On day 4, each of the 6 independent lineages was constructed by picking two N2 L4 hermaphrodite worms from the initial F0 expanded population to a new plate to produce the subsequent generation (generation 2). On day 5, embryos were collected from the original plate (generation 0). This procedure was repeated to propagate the lineages for 20 subsequent generations. Embryos were collected for every two subsequent generations (0, 2, 4, 6, 8, 10, 12, 14, 16, 18 and 20).

### Embryo collection

To collect the embryos, both worms and embryos were washed off plates into 15 ml falcon tubes with 0.1% Triton X. The plates were washed several times to maximise the collection of worms and embryos. The falcon tubes were centrifuged at 2000 RPM for 1 minute and the supernatant was removed. Remaining embryos on the washed plates were loosened from agar by spraying 0.1% Triton X with the pipette angled against the plate. The volume of liquid containing additional washed off embryos was added to the worm pellet. Three further washes of the pellet were performed until the supernatant was clear to ensure that OP50 bacteria had been removed. Bleaching procedure was used to retain only the embryonic material: bleaching process destroys the worms while leaving the embryos untouched. Worms were bleached using hypochlorite treatment in which 5.5 ml of bleach was added to each sample. Samples were vortexed continuously for 5 minutes with intermittent checking to ensure worm carcasses were dissolving and to avoid over exposure of embryos to bleach. Bleach solution was washed out rapidly once worm carcasses had disappeared with addition of M9 to dilute bleach, centrifugation to pellet undissolved embryo material, complete removal of supernatant and repeating this procedure for a total of three washes. Following the 3 post bleaching washes, the supernatant was removed from the embryo pellet. The pellet was resuspended in 1 ml 0.1% Triton X and then split into 1.5ml microcentrifuge tubes follows: 25% of each sample was reserved for RNA library preparation and 100ul TRIzol was added to inhibit RNase and maintain RNA integrity. 75% of each sample was reserved for ATAC-seq and frozen with no additive. Microcentrifuge tubes were then submerged in liquid nitrogen before being stored at – 80 □C.

### Extraction of material for assessment of epimutations

#### RNA extraction

RNA was extracted via chloroform Isopropanol extraction following standard protocols which had been optimised for our lab^9, 25^. In brief, embryos in TRIzol were lysed through 5 freeze-cracking cycles (frozen in liquid nitrogen and then thawed in a 37 □C water bath). Tubes containing lysed embryos were vortexed for 5 minutes with 30 second pauses at 30 second intervals, and then incubated at room temperature for 5 minutes for disruption of RNA-protein complexes. 200μl chloroform per ml of TRIzol were added to each sample followed by vigorous shaking. Samples were then incubated for two to three minutes at room temperature followed by centrifugation at a maximum speed at 4 □C for 10 minutes. The top aqueous layer was aspirated and transferred to a new tube into which 1μl glycogen, and an equal volume of isopropanol was added. RNA was precipitated overnight at -20 □C. After overnight precipitation, the samples were centrifuged for 1 hour at 4 □C. The supernatant was removed and 500μl of 75% ETOH were added to the pellet. The samples were centrifuged at max speed for 10 minutes at 4 □C. All ETOH was removed, and the pellet allowed to air dry. The RNA pellet was resuspended in 10μl H20. RNA sample concentration and quality were quantified on a 2200 TapeStation instrument using Agilent RNA screen tapes. An RNA Intengrity (RIN) score was derived. Samples were additionally quantified using Nanodrop.

#### Small RNA extraction

As previously described^11^, the TruSeq Small RNA Prep Kit (Illumina) was used to prepare small RNA libraries from RNA pyrophosphohydrolase (Rpph) treated RNA. RNA was mixed with 1.5μl RPPH enzyme and 2μl 10X NEB Buffer 2 in a final volume of 20μl. This was incubated at 37 □C for 1 hour. The number of PCR cycles was increased from 11 to 15 as per protocol optimised previously^9^, otherwise all steps were done according to manufacturer’s instructions. Size selection was performed using gel extraction with a 6% TBE gel (Invitrogen). Validation of 22G-RNA libraries was done on a 2200 TapeStation (Agilent).

#### Assay for Tranposase Accessible Chromatin (ATAC)

The protocol for ATAC was adapted from Daugherty^26^ after the original method for ATAC-seq^27^ and was previously described^11^. In brief, nuclei were extracted from embryo samples through three cycles of freeze cracking. 200μl of Nuclear Preparation Buffer added to each microcentrifuge tube containing frozen embryos, sample submersion in liquid nitrogen for 90 seconds and then transferred to a 37 □C water bath for 90 seconds. Embryonic material was then transferred to a Wheaton glass tissue homogenizer in ice and was ground (3.5 grinds), vessels containing the embryonic material were covered and centrifuged for two minutes at 200 RCF at 4 □C. The supernatant containing nuclei was transferred to a microcentrifuge tube on ice. The remaining embryo material was ground again, spun down and the supernatant again transferred to the collection tube. This cycle was repeated 4 times. Collection tubes were spun at 1000 RCF at 4 □C for 10 minutes, supernatant was discarded, leaving the nuclei pellet behind. To each sample nuclei pellet, Tagmentation enzyme and Tagmentation buffer were added (Illumina Tagment DNA enzyme and buffer). The samples were incubated for 30 minutes at 37 □C on a Thermoshaker set to 580 RPM. Samples then underwent DNA clean up with a Qiagen MiniElute Reaction Cleanup Kit. The resulting DNA was eluted in 10μl of EB buffer. PCR adapters from the Illumina Nextera DNA prep kit (Illumina) were selected and added to each sample along with PCR master mix. PCR was run with the following cycle parameters: 72 □C for 5 minutes, 98 □C for 30 seconds, 14 cycles (98 □C for 10 seconds, 63 □C for 30 seconds, 72 □C for 1 minute). 4 □C hold. Size selection isolates the desired DNA fragments and was achieved with magnetic AMP X beads (Beckman Coulter). First, excessively large DNA fragments were removed as follows; 25μl AMP X bead solution was added to each sample. Samples were incubated at room temperature for 10 minutes. The PCR tubes containing the bead and sample mix were then put onto a magnetic rack for 5 minutes until the sample appeared clear. The samples were transferred without disturbing the magnetic beads to a new set of PCR tubes. To remove excessively small DNA fragments, the same procedure was followed, but the volume of AMP X bead solution added was adjusted to 65μl. After incubating on the magnetic rack, the clear liquid was carefully removed and discarded. At this point, the beads had the desired library bound. The bead pellet was washed twice with 80% EtOH and then allowed to air dry at room temperature for two minutes. Excess EtOH droplets were removed with a pipette tip from inside each PCR tube. 22μl of nuclease free water was added to each sample and the beads were washed down and dispersed into the liquid. The PCR tubes were returned to the magnetic rack, the DNA having eluted into the nuclease free water. The liquid was removed without disturbing the magnetic beads to a final collection tube which could then be frozen at -80 □C. Validation of ATAC libraries was performed by quantification on the 2200 TapeStation using Agilent D1000 Screen tapes. ATAC libraries were also quantified on Qubit using the Qubit dsDNA HS Assay kit from Invitrogen.

### Library construction

#### RNA library construction

cDNA library preparation and PolyA-selected RNA sequencing were carried out in the MRC LMS Genomics Facility to generate PE100bp reads. A cDNA library from 31 samples was prepared using the TruSeq RNA Sample Prep kit (Illumina) following the manufacturer’s instructions. The cDNA library was then hybridised to the flow cell of the Illumina HiSeq 2500 and sequenced. The library was run on 3 separate lanes to achieve good coverage depth. The sequencing data were processed by the instrument’s Real Time Analysis (RTA) software application, version 1.18.64. De-multiplexing was done within the sequencing facility with CASAVA.

#### ATAC library construction

For submission to LMS Genomics Facility, multiplexing of samples was performed, with 10–12 samples grouped into a pool. Samples were combined as 10nM dilutions. Paired end sequencing of ATAC samples was done in the LMS Genomics Facility on a HiSeq instrument. The sequencing data were processed by the instrument’s Real Time Analysis (RTA) software application, version 1.18.66.3 using default filter and quality settings. Demultiplexing was done within the sequencing facility using CASAVA-2.17, allowing zero mismatches.

#### Small RNA library construction

For submission to LMS Genomics Facility, 3nM dilutions were prepared from each small RNA sample. 3nM sample dilutions were pooled at ∼ 30 samples per pool with unique indexes. Sample pools were validated on TapeStation and quantified on Qubit using dsDNA reagents. MiSEQ Single Read sequencing (50 bp read length) was used to validate sample pools. Balancing or preparation of new pools was undertaken to improve the balance of samples in the pool as needed. High Output single read sequencing was performed on NExtSeq 2000 instrument in the LMS Genomics Facility. The max read length was 60 bpm. The sequencing data were processed by the instrument’s Real Time Analysis (RTA) software application, version 2.11.3.

### Pre-processing and alignments

#### RNA-seq libraries

RNA fastq files were aligned to the *C. elegans* genome using Bowtie2^28^ to produce a file containing the raw counts data. Sam files were sorted and converted to bam using samtools^29^ and then bed files using bedtools^30^. Sorted counts were intersected with coordinates of coding genes derived from the UCSC genome browser^31^using BEDTools intersect commands^30^:

The files were simplified so that only the longest transcript was represented (the other isoforms were filtered out) and 21U-RNAs were removed. In addition, all genes for which no expression was seen in any of the samples were removed. The read values were normalised using DESeq2^32^.

#### Small RNA

Secondary processing and demultiplexing of data were done within the sequencing facility with Bcl2fastq 2_2.20.0 (Illumina). Identification of the small RNAs was further done using the tinyRNA pipeline (tinyRNA-1.0.1)^33^ following the standard parameters. Only the following options have been adapted: adapter sequence to trim = AGATCGGAAGAGCACACGTCTGAACTCCAGTCAC; Trim bases from the tail of a read = 15. Read counts were then normalized using DESeq2^32^.

#### ATAC-seq libraries

Trimming of reads to remove adapters was performed using FASTX Toolkit from the Hannon Lab (RRID:SCR_005534). Sequence alignment was carried out using Bowtie2^28^ to produce a file containing the raw counts data. Sam files were sorted and converted to bam using samtools^29^.

### Identification of gene expression changes

Global differentially expressed genes (DEGs) using generations as replicates were identified with DESeq2^32^ by comparing low dose and control conditions and high dose and control conditions. Specific gene expression changes observed in each generation were detected using the DESeq normalized count matrix. Gene expression changes were identified using a local regression model (loess) in which log_2_(mean counts) for each generation (1 to 20) were plotted against the log_2_(mean counts) of the generation 0 (pre-accumulation line) for each gene in line with the approach taken in previous work to identify changes in RNAs^9^. For each data point, a Z-score was derived by subtracting the mean of the residuals from the linear model from each individual residual value and dividing the output by the standard deviation of the residuals. Regions with a Z-score greater or less than 2.25 were defined as showing differential accessibility. Regions with a Z-score greater than 2.25 were annotated as ‘Up’ epimutations and regions with a Z-score less than 2.25 were annotated as ‘Down’ epimutations. The Z-scores obtained for expression changes and epimutations were cut offs and used to get binarized tables (z-scores < -2.25 set up as “-1” (down epimutation); -2.25 < z-score < 2.25 as “0” and z-scores > 2.25 as “1” (up epimutation)) for each locus as previously described^11^.

### Identification of genetic mutations

ATAC-seq data were used to identify SNPs and indels. Briefly, the *C. elegans* genome (version WS252) was indexed using samtools^29^ and the following command: *samtools index genome.fa*. Bam files from the ATAC-seq libraries preparation were used to generate pileup files using samtools^29^ and VarScan (v2.4.5)^34^was used to detect DNA mutations in each sample.

In each lineage, DNA mutations detected in F0 were used as a baseline and discarded from each subsequent generational dataset. Then, true, and consistent mutations were separated from artefacts by keeping only the mutations that appeared in each of the generations and remained until the last generation. As with identified a lot of false positive (mutations that appeared at one generation but didn’t remain in the successive ones) we kept data only for the generations that could be verified using the datasets of the following generations. Therefore, the new DNA mutations appearing only in the last generations available (generation 20 for control lineages, 16 for LD lineages, and 18 for HD lineages) were discarded from the analysis. True DNA mutations were identified in each generation and each lineages using a customized R script. Tables were computed for SNPs and indels with the positions of the mutations for each generation. Sum up tables were then manually constructed with the total number of genes affected by DNA mutations, the generation to which the epimutation appears, the nucleotide of reference and the new one after SNP. Reference codons and new ones after indels were recorded using the jbrowser (Wormbase) and the input from Varscan.

### Identification of small RNAs epimutations

The feature counts table computed by the tinyRNA pipeline was used as input for subsequent analysis on Rstudio (R version 4.2.1). As for the gene expression changes identification, after the normalisation step of count data using DESeq2, epimutations were identified using a linear model in which log_2_(mean counts) for each generation (1 to 20) were plotted against the log_2_(mean counts) of the generation 0 (pre-accumulation line) for each gene in line with the approach taken in previous work to identify epimutations in small RNAs^9^. For each data point, a Z-score was derived by subtracting the mean of the residuals from the linear model from each individual residual value and dividing the output by the standard deviation of the residuals. Regions with a Z-score greater or less than 2.25 were defined as showing differential accessibility. Regions with a Z-score greater than 2.25 were annotated as ‘Up’ epimutations and regions with a Z-score less than 2.25 were annotated as ‘Down’ epimutations. The Z-scores obtained for expression changes and epimutations were cut offs and used to get binarized tables (z-scores < -2.25 set up as “-1” (down epimutation); -2.25 < z-score < 2.25 as “0” and z-scores > 2.25 as “1” (up epimutation)) for each locus as previously described^11^.

Data from all sRNAs were later sorted by type of interest: 22G-RNAs, 26G-RNAs, piRNAs, and miRNAs using the value or tag identified by the tinyRNA pipeline (Sup. Table 1). tRNAs were sorted among the unknown (“unk” in Sup. Table 1) reads from the tinyRNA pipeline using the reference annotation WS279 (WormBase Release WS279_master.gff3).

**Table 1:** Genes showing both a change in expression level and a DNA mutation. WB = WormBase identification number. Gen. exp chg. = Generations with gene expression change. +/- column indicates if the observed change was overexpression (+) or underepression (-). Gen. mut. = Generations showing DNA mutation. Data were missing for DNA mutation identification at the generations 18 and 20 for Low dose L2 but in the gene Y43D4A.6, the SNP appeared at the generation 10 and up to the generation 16 we thus assumed that it would also be observed in the subsequent generations if it’s a true mutation.

| Condition | Lineage | Gene (WB) | Gene | Gen. exp. chg. | +/- | Gen. mut | Mut. type |
| --- | --- | --- | --- | --- | --- | --- | --- |
| Low dose | L1 | WBGene00019132 | bath-19 | 2_10_16 | + | 16 | Indel |
| Low dose | L1 | WBGene00018024 | F35A5.1 | 2_10_16 | + | 16 | SNP |
| Low dose | L2 | WBGene00012792 | Y43D4A.6 | 20 | + | 20? | SNP |
| High dose | H2 | WBGene00001839 | hdl-1 | 18 | - | 18 | SNP |

### Test for gene expression changes/epimutation inheritance

All our datasets derived from a pre-mutation founder generation and comes from alternate embryos sampling realized every two generations (2, 4, 6, 8, 10, 12, 14, 16, 18 and 20). We had to make certain assumptions about the epimutation status for the missing generations and considered for instance that an epimutation appearing in the generation 2 and present in the generation 4 would exist in the generation 3 (even if data were not collected at that generation). Thus, in order to be considered to be inherited, we required that two consecutive even numbered generations displayed an epimutation. Moreover, due to insufficient yield of RNA or cDNA some samples were omitted from sequencing (RNA-seq dataset generation 12 for the line C2 and generation 20 for the line H1; small RNA-seq dataset generation 6 for the lines C2, L2 and H1, generation 10 for the line L1; ATAC-seq dataset generations 8, 10 and 14 for the line C2, generations 6, 14, 18 and 20 for the line L1, generations 2, 6, 14,18 and 20 for the line L2, generations 12, 14, 16 and 20 for the line H1 and generations 6 and 20 for the line H2).

In addition, each change of directionality in an epimutation from a generation to the subsequent one (for instance “Up” epimutation at generation 2 and “Down” at generation 4) was considered to be a termination of the preceding epimutation and commencement of a new epimutation run.

### Characterisation of tRNA fragments in epimutation accumulation lines

For further characterization, *C. elegans* tRNAs were downloaded from http://gtrnadb.ucsc.edu. The sequences were then aligned to the genome using Bowtie^28^ with 0 mismatches. The resulting sam file was converted to bam using samtools^29^ and bed using bedtools^30^ to give coordinates of tRNA genes. To assign short non-coding RNAs to tRNAs, alignment files for 18-36 nucleotide small RNAs generated by tinyRNA (above) were converted to bed files using bedtools and then intersected with the tRNA coordinates using bedtools intersect. The position of the start of the read relative to the tRNA sequence was obtained from this file using R^35^. Readcount for each small RNA mapping to a tRNA was normalized using the size factors from DESeq calculated as part of the tinyRNA pipeline. Counts for individual tRNA types were obtained by summing normalized reads mapping to the 3’ half of each tRNA type and this table (Sup. Table. 2) was subsequently used for further analysis to identify epimutations in tRNAs according to the procedures in section “*Identification of small RNAs epimutations*” above. To identify potential target genes, small RNAs corresponding to tRNA 3’ halves were extracted from all the lines and combined into a single file, along with the annotation indicating which tRNA the sequence was derived from. This was aligned to the ce11 genome allowing up to 3 mismatches. Protein-coding genes overlapping with mismatched tRNA fragments were then obtained using bedtools intersect, to identify genes that were potentially targeted by tRNAs (Sup. Table. 3). Code for this analysis and the figures is on the GitHub page.

### Identification of Argonautes that bind tRNAs

Fastq files corresponding to sequences immunoprecipitated in association with *C. elegans* Argonautes were downloaded from the SRA via GSE208702^18^. Fastq files were converted to fasta files and aligned to ce11 using bowtie^28^ with 0 mismatches and recording only one match per sequence. The resulting sam files were converted to bam using samtools^29^ and bed files using bedtools^30^. These files were intersected with the tRNA annotations as above and reads greater than 25 nucleotides were selected to ensure that no overlapping 22G-RNAs were included. Reads were normalized to the total read count, and further normalized by the paired input for each IP. Total tRNA normalized tRNA counts were then visualised to identify which AGOs appeared to show notable binding. To test whether the identified Argonautes were required to stabilise tRNAs, small RNA reads were downloaded from mutants lacking ergo-1, wago-10 and rde-1 as well as N2 (WT) and reads were assigned to tRNA fragments as before, again ensuring that 22G-RNAs were not included. Reads were normalized to total read count. Total tRNA counts were compared for the mutants. The apparent decrease in wago-10 mutants was then explored further by investigating counts mapping to different tRNA types across the three replicates, using a Wilcox unpaired test to identify any that showed significant differences between N2 and wago10 mutants. All code for the R analysis including figure plotting commands is on the GitHub page.

### Integration of gene expression change and epimutations data

To match up gene expression changes and sRNAs epimutations, we integrated data in a gene-centric manner. For each gene, we recorded its expression state, if expression state change was inherited and the presence of epimutated sRNAs. We also recorded the directionality of each change (Up or Down gene expression change or epimutation) and the generations affected by both the gene expression change and the epimutation to investigate simultaneous (at the same time) and concordant (in the same direction) changes.

### Gene ontology analysis

We computed lists of genes with expression changes or epimutation and uploaded them to WormEnrichr^36, 37^. The number of genes annotated for each identified ontology term was compared between the test lists and the background sets of genes (with no epimutations or expression changes). We set up as a requirement a minimum of 5 genes to be present in either the test or background sample gene list for the ontology term to be included in the analysis. Fisher’s Exact test was used to calculate the enrichment of test list of gene for ontology terms in comparison to the background lists.

### Production of plots and figures

All plots and statistical analysis were made using R^35^ and RStudio (RStudio Team (2022). RStudio: Integrated Development Environment for R. RStudio, PBC, Boston, MA URL. http://www.rstudio.com/. All code for the R analysis including figure plotting commands is on the GitHub page.

Plots and figures were then processed with Affinity Designer 2.

## Results

### 1. Chronic Cisplatin exposure induces genotoxic stress and gene expression changes

To investigate how epimutations might be affected by genotoxic stress we established parallel lines of *C. elegans*, propagated under minimal selection, in control conditions, and exposed to either 75uM or 150uM cisplatin (henceforth low and high dose), selected such that the treatment had a notable effect on embryonic lethality but did not block propagation (Sup. Fig. 1.A). High dose cisplatin led to a significant increase in the rate of indels (Fig. 1. A), confirming that the treatment induced genotoxic stress, as expected based on the known mechanism of cisplatin in inducing DNA damage through inter and intrastrand cross links^38^. Nevertheless, the total number of mutations induced over the course of the experiment was still relatively small (at around 13.5 snps and 7 indels by 20 generations), thus the rate of mutation even under cisplatin exposure is still orders of magnitude lower than the expected rate of epimutations^9, 11^.

**Figure 1:**
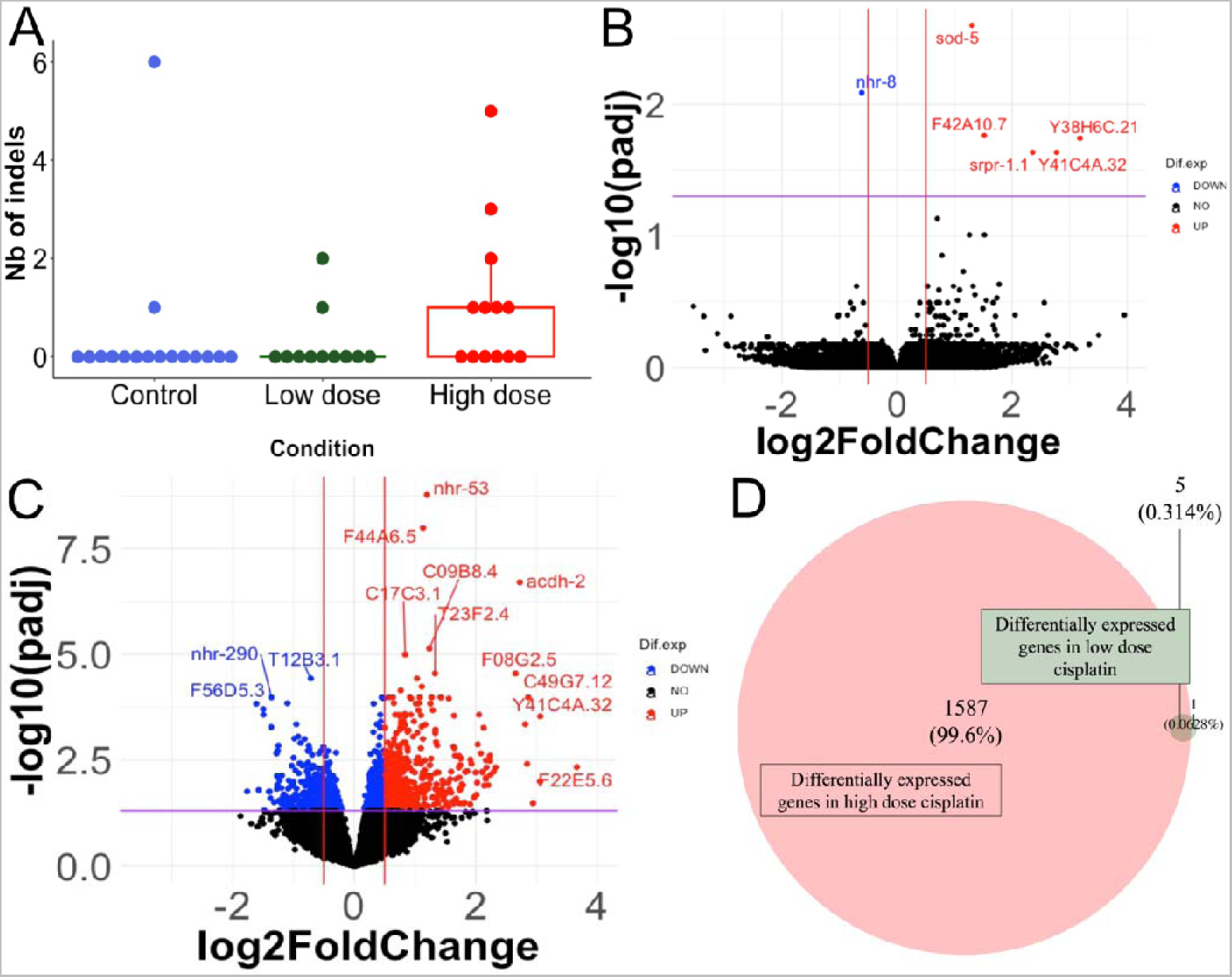
Effect of cisplatin on gene expression and DNA mutations. A. Boxplot of the number of indels arising in each generation for control condition (blue), cisplatin low dose (green) and cisplatin high dose (red). A significant increase in indels number was observed in high dose cisplatin (Kruskal-Wallis rank-sum test (p-value= 0.04642) followed by Conover-Iman test: Control vs. High dose, p-value= 0.0090; Control vs. Low dose, p-value= 0.3972; Low dose vs. High dose, p-value= 0.0260) B. Volcano plot representing gene expression changes in low dose cisplatin compared to control. X-axis = log2FoldChange of the genes, y-axis = -log10(adjusted p-value). Both Foldchange values and adjusted p-values were calculated using DESeq2. Genes in red are significantly up-regulated, genes in blue are significantly down-regulated and genes in black showed no change in expression. C. Volcano plot representing gene expression changes in high dose cisplatin compared to control. X-axis = log2FoldChange of the genes, y-axis = -log10(adjusted p-value). Both Foldchange values and adjusted p-values were calculated using DESeq2. Genes in red are significantly up-regulated, genes in blue are significantly down-regulated and genes in black showed no change in expression. D. Venn diagram of differentially expressed genes in high dose cisplatin compared to control (red), differentially expressed genes in low dose cisplatin compared to control and genes differentially expressed in both condition (low and high dose). 6 genes were differentially expressed in low dose condition, 1592 in high dose and 5 genes shared differentially expression in the two conditions.

Chronic exposure to cisplatin could induce gene expression changes directly, independent of epimutations. To identify these effects, we looked for genes that showed consistent changes in expression across all generations of the experiment. We identified 6 differentially expressed (DE) genes in low dose compared to control (Fig.1.B) and 1 592 DE genes in high dose compared to control (DESeq2, Wald-test, p-value< 0.05) (Fig.1.C). 5 DE genes were shared between low and high dose conditions (Fig.1.D). Since only a few DE genes were observed in low dose, no relevant enriched function was identified during the GO term enrichment analysis. However, 279 significantly enriched functions in high dose cisplatin condition compared to control were observed (Sup. Table. 4). The gene expression differences observed support the conclusion that cisplatin exposure induces genotoxic stress (Sup. Fig. 1.B).

We next investigated the potential of DNA mutations to be responsible for gene expression changes in our dataset. We examined changes in gene expression that occurred relative to the first generation of each lineage, where changes in expression were unlikely to be caused by exposure to cisplatin. We identified genes showing both expression change and DNA mutation at the same generation (Table 1). No genes underwent both kind of changes in the control condition. In the cisplatin low dose condition, three genes had both expression changes and DNA mutation and only one gene in cisplatin high dose condition. Thus, only a small number of genes with expression changes over the course of the experiment were associated with DNA mutations (Control: 0/11497; Low dose: 3/10058; High dose: 1/9737). In addition, the two genes in L1 lineage with both change in expression and DNA mutation had a change in expression at three different generations and a DNA mutation observed in only one of them, making a correlation between the DNA mutation and the change in expression rather unlikely.

In conclusion, the cisplatin treatment did impact the rate of DNA mutations with a correlation between an increase in dose and an increase in the number of new mutations observed. This result is not surprising as cisplatin is a genotoxic compound and exposure to cisplatin had led to DNA mutational signature in several organisms^39–41^. Despite the increase in the rate of DNA mutations, a correlation between DNA mutations and gene expression changes concerned only a very limited number of genes.

### 2. Cisplatin exposure has limited effect on the rate and spectrum of heritable gene expression changes

Having established that cisplatin exposure induced genotoxic stress, we sought to decipher whether the properties of epimutations were affected. Importantly, we wanted to specifically identify epimutations rather than direct transcriptional responses to cisplatin. We therefore investigated changes in gene expression that occurred in each individual generations, within the individual lineages and within the time course of the experiment.

We first observed that the number of new epimutations affecting gene expression was not significantly different between control and cisplatin-treated lines (Kruskal-Wallis rank sum test, p-value = 0.112) (Fig.2.A). The duration of epimutations affecting gene expression showed a significant difference across conditions (Log-rank test, p-value < 0.0001). Epimutations tended to last slightly longer under conditions of high dose cisplatin, and slightly shorter in low dose cisplatin relative to control conditions (Fig.2.B). However, these differences were small, as demonstrated by an analysis of the fraction of changes in expression seen each generation that were inherited, which was similar between all conditions (Fig.2.C).

**Figure 2:**
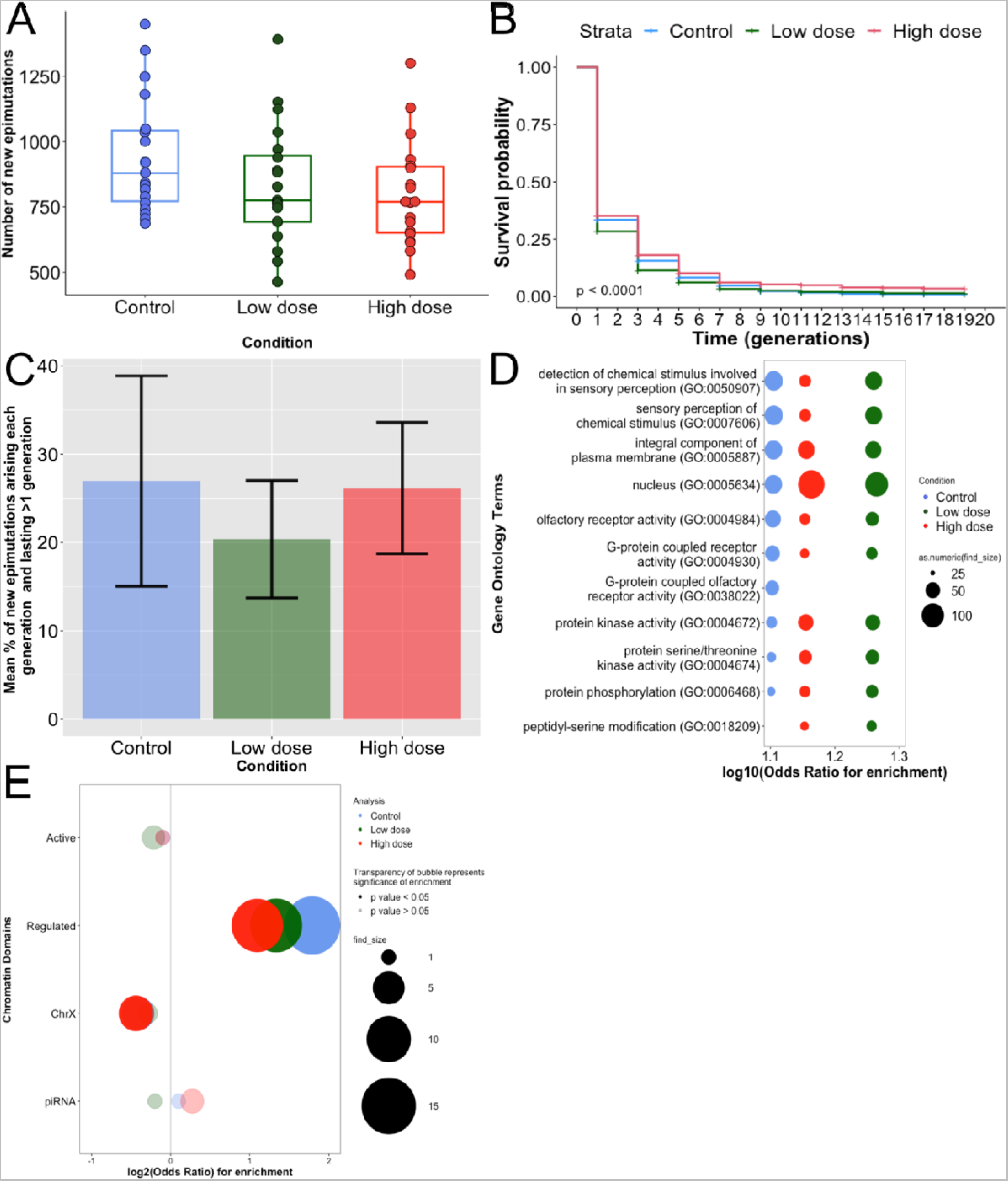
Effects of cisplatin exposure on gene expression epimutations. A. Boxplot of the number of new RNA epimutations arising at each generation of the MA lines compared to the pre-mutation generation F0 and for each condition: control (blue), cisplatin low dose (green) and cisplatin high dose (red). No significant difference was observed between the conditions (Kruskal-Wallis rank-sum test, p-value= 0.112). B. Survival curves representing the new RNA epimutations duration in each exposure conditions: control (blue), cisplatin low dose (green) and cisplatin high dose (red). We observed a significant difference between the three conditions with an increased duration of epimutations in high dose and a decreased duration in low dose compared to control (Log-rank test, p-value < 0.0001). C. Barplot of the mean percentage of new RNA epimutations that lasted more than 1 generation compared to total epimutations arising each generation. Data are presented by condition: control (blue), cisplatin low dose (green) and cisplatin high dose (red). The mean percentages of lasting epimutations compared to all epimutations were 26.9% for control, 20.4% for low dose and 26.2% for high dose. No significant difference was observed between conditions. D. Bubble plot illustrating ontology term enrichment of RNA epimutations lasting more than 1 generation in control (blue), cisplatin low dose (green) and cisplatin high dose (red) compared to gene without epimutation. Enrichment was calculated with _χ_-squared test. The top 10 results per condition are shown. X-axis shows log10(_χ_) for enrichment. Y-axis shows ontology terms. All displayed ontology terms were significantly enriched. E. Bubble plot showing the distribution of lasting RNA epimutations in control (blue), cisplatin low dose (green) and cisplatin high dose (red). Y-axis displays the constitutive chromatin domains investigated. X-axis shows the log2(Odds) of enrichment. Odds ratio and p-values were calculated using Fisher’s Exact Test with Bonferroni correction. p-value cut off for significance is 0.05.

We next investigated whether the spectrum of genes affected by epimutations was modified by cisplatin exposure. Using gene ontology analysis, we identified some significantly enriched GO terms that were specific to cisplatin exposure (Sup. Table. 5); however, there was a strong overlap between the top 10 enriched GO terms in all conditions (Fig.2.D), indicating that cisplatin exposure did not result in large shifts in genes affected by epimutations.

We also looked at the chromatin environment of genes affected by epimutations (Fig.2.E). Under control conditions, epimutations were enriched in regions rich in H3K27me3 and depleted from active regions enriched in H3K36me3, and from the X chromosome which was consistent with our previous results^11^. Cisplatin exposure did not alter these trends, supporting the conclusion that cisplatin exposure did not change the spectrum of epimutations.

### 3. Cisplatin exposure does not affect the rate or spectrum of sncRNAs-mediated epimutations

Small non-coding RNAs are well established as carriers of transgenerational epigenetic inheritance in *C. elega*ns^17^. We first investigated 22G-RNAs, as they are the major source of epimutations, and found no difference between control and cisplatin-treated lines in term of epimutation rate (Fig.3.A) nor a cisplatin dose-dependent change of epimutations duration (Fig.3.B).

**Figure 3:**
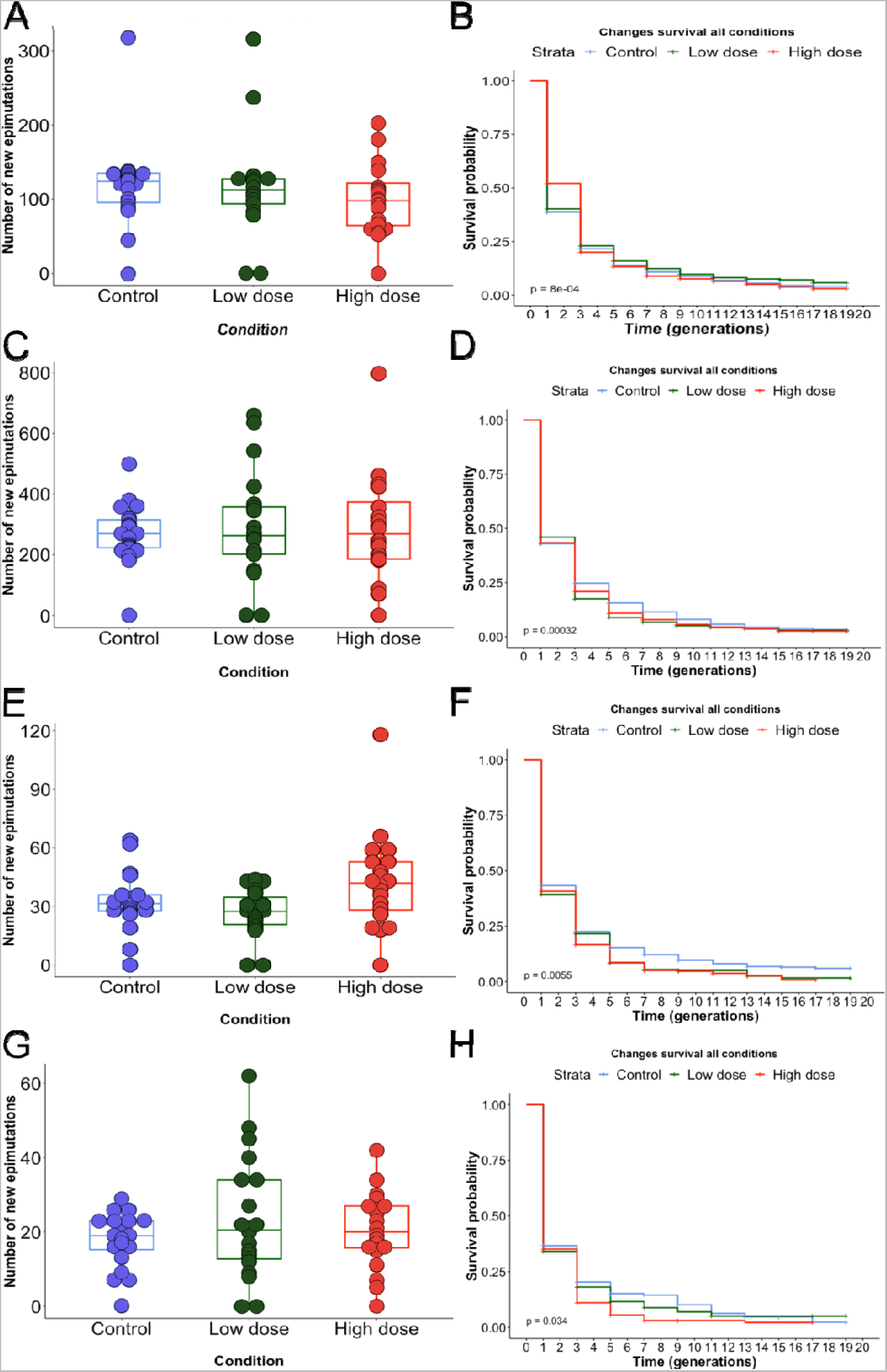
Effects of cisplatin exposure on sncRNAs epimutations. A. Boxplot of 22G-RNAs epimutations rate for each generation of the MA lines compared to the pre-mutation generation F0 and for each condition: control (blue), cisplatin low dose (green) and cisplatin high dose (red). No significant difference was observed between conditions (Kruskal-Wallis rank-sum test, p-value = 0.3934). B. Survival curves representing the new 22G-RNAs epimutations duration in each exposure conditions: control (blue), cisplatin low dose (green) and cisplatin high dose (red). We observed a significant difference between the three conditions (Log-rank test, p-value = 8e-04) but the difference wasn’t dose dependent. C. Boxplot of piRNAs epimutation rate for each generation of the MA lines compared to the pre-mutation generation F0 and for each condition: control (blue), cisplatin low dose (green) and cisplatin high dose (red). No significant difference was observed between the conditions (Kruskal-Wallis rank-sum test, p-value= 0.9901). D. Survival curves representing the new piRNAs epimutations duration in each exposure conditions: control (blue), cisplatin low dose (green) and cisplatin high dose (red). We observed a significant difference between the three conditions (Log-rank test, p-value = 0.00032) with slightly higher survival changes of epimutation in control condition. E. Boxplot of miRNAs epimutations rate for each generation of the MA lines compared to the pre-mutation generation F0 and for each condition: control (blue), cisplatin low dose (green) and cisplatin high dose (red). No significant difference was observed between conditions (Dunn Kruskal-Wallis multiple comparison, Control vs. LD: p-value = 0.22979622; Control vs. HD: p-value= 0.32832185). F. Survival curves representing the new miRNAs epimutations duration in each exposure conditions: control (blue), cisplatin low dose (green) and cisplatin high dose (red). We observed a significant difference between the three conditions (Log-rank test, p-value = 0.0055) with slightly higher survival changes of epimutation in control condition. G. Boxplot of 26G-RNAs epimutations rate for each generation of the MA lines compared to the pre-mutation generation F0 and for each condition: control (blue), cisplatin low dose (green) and cisplatin high dose (red). No significant difference was observed between conditions (Kruskal-Wallis rank-sum test, p-value = 0.7074). H. Survival curves representing the new 26G-RNAs epimutations duration in each exposure conditions: control (blue), cisplatin low dose (green) and cisplatin high dose (red). We observed a significant difference between the three conditions (Log-rank test, p-value = 0.034) with slightly higher survival changes of epimutation in cisplatin low dose condition.

We previously found that 22G-RNAs epimutations were overlapping with changes in gene expression and genes showing heritable changes in gene expression were significantly enriched for simultaneous 22G-RNA-based epimutations, especially for genes linked to xenobiotic defence^11^. Therefore, we investigated the association between 22G-RNAs and gene expression changes in our control and cisplatin-treated lines. Amongst genes that had inherited RNA expression changes and were targeted by 22G-RNAs epimutations, we calculated the percentage of which that had simultaneous inherited 22G-RNAs epimutations, the percentage of which that had non-inherited 22G-RNAs epimutations and the percentage of which that had non-simultaneous 22G-RNAs epimutations, and thus for each condition (Sup. Fig. 2. A. left stacked-barplots in each panel). We did the same among the genes that had non-inherited RNA expression changes and were targeted by 22G-RNAs epimutations (Sup. Fig. 2. A. right stacked-barplots in each panel). We found quite similar results for both control and low dose cisplatin conditions (37.5% of genes with inherited expression change and targeted 22G-RNA epimutation had a simultaneous and inherited 22G-RNA epimutations in control, 33.3% in low dose) with and a decrease in the percentage of genes (20%) with both simultaneous and inherited expression changes and 22G-RNAs epimutations for cisplatin high dose condition.

Previously we demonstrated that gene expression changes that were inherited transgenerationally were more likely to be associated with epimutations in 22G-RNAs. We recapitulated this observation in control, low dose and high dose cisplatin using an association analysis. However, the enrichments were reduced in high dose cisplatin, suggesting that the association between 22G-RNA changes and gene expression changes was weakened by the exposure to cisplatin (Sup. Table 6).

Our next step was to look at piRNAs; involved in transgenerational epigenetic inheritance in *C. elegans*^42^, and miRNAs that have been suspected to play a role in epigenetic inheritance in medaka fish^43^ or in rat^44^ after exposure to chemicals. In addition, we had found in a previous study that both miRNAs and piRNAs could be inherited transgenerationally^11^. Results revealed no significant difference due to the cisplatin treatments for both piRNAs epimutation rate (Fig.3.C) and epimutation duration (Fig.3.D). Similarly, we observed no difference in the total number of new miRNAs epimutations per generation between control and treated lines (Fig.3.E) or in the duration of epimutations (Fig.3.F).

Finally, we investigated 26G-RNAs that are small interfering RNAs (siRNAs) enriched in the germline and involved in the 22G-RNAs synthesis^45^. Once again, no overall significant difference in epimutations rate (Fig.3.G) or the epimutations duration (Fig.3.H) between conditions was identified.

### 4. Cisplatin exposure affects the rate of tRNAs epimutations

In the course of our analyses of sncRNAs we noticed that there appeared to be more reads mapping to tRNA loci under cisplatin conditions (Fig.4.A). We were intrigued by this because tRNA halves are known to be abundant in mammalian cells and have been implicated in intergenerational epigenetic transmission through mammalian sperm^4, 44, 46–48^. We therefore decided to investigate whether tRNA might be induced by cisplatin treatment. We mapped sncRNAs to tRNAs, demonstrating that the predominant initiation point for sncRNAs coincided with the 3’ half of the tRNA (Fig.4.B). Taken across all generations there was a significant increase in the reads mapping specifically to the 3’ half of the tRNA in both low and high dose cisplatin, and a trend towards increased reads in high dose cisplatin (p<2e-16, Jonckhere test for ordered medians, Fig.4.C). The most abundant tRNAs fragments in all conditions combined were GlycineGCC, GlutamineTTG and Glutamic acidCTC (Fig.4.D). 3 tRNAs showed a significant increase in expression in cisplatin high dose: GlutamineTTG, ValineAAC and SerineCGA (DESeq2: adjusted p-value < 0.05, Fig.4.E).

**Figure 4:**
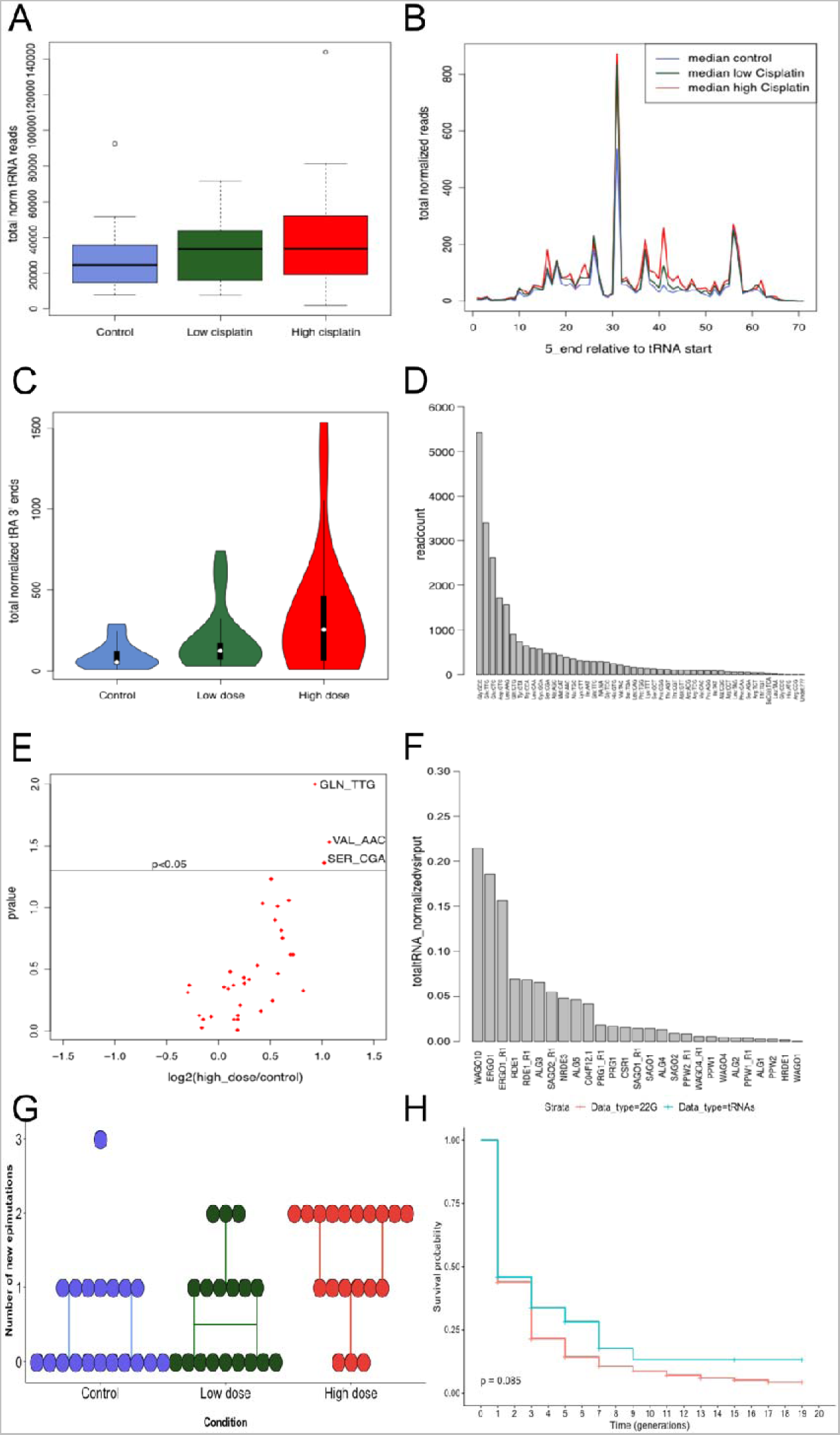
Cisplatin effects on tRNAs. A. Total normalized reads that mapped to tRNAs in each condition: control condition (blue), cisplatin low dose condition (green) and cisplatin high dose condition (red). Total normalized reads represented according to their mapping positions on the respective tRNA sequence and in each different condition: control condition (blue), cisplatin low dose condition (green) and cisplatin high dose condition (red). C. Violin plot of total normalized tRNAs mapping specifically to the 3’ half of the tRNAs in the different conditions: control condition (blue), cisplatin low dose condition (green) and cisplatin high dose condition (red). A significant increase was observed in both low and high dose cisplatin, with a trend towards increased reads in high dose cisplatin (p<2e-16, Jonckheere test for ordered medians. D. Bar plot showing the number of read counts associated to each kind of tRNAs. The tRNAs with the most associated fragments were GlycineGCC, GlutamineTTG and Glutamic acidCTC. E. Volcano plot of expression change between high dose and control (in log2(fold change)) for the different kind of tRNAs. Points above the p<0.05 line show significant expression change for this corresponding tRNAs. Adjusted p-values were calculated using DESeq2. F. Bar plot of the total normalized 3’ halves tRNAs reads and their association to each AGO protein. Both Ergo1 and Wago10 showed a high enrichment in tRNA 3’ halves relative to control Ips. G. Boxplot of the number of new epimutations affecting tRNAs fragments arising at each generation of the MA lines compared to the pre-mutation generation F0 and for each condition: control (blue), cisplatin low dose (green) and cisplatin high dose (red). A significant increase was observed in cisplatin high dose condition (Kruskal-Wallis rank sum test followed by pairwise Wilcoxon test with Bonferroni correction: HD vs. C: p-value = 0.003649917, HD vs. LD: p-value = 0.020226741). H. Survival curves representing the duration of epimutations in 22G-RNAs (red) and 3’halves tRNAs fragments (blue). No significant difference was observed between the two types of snRNAs (Log-rank test, p-value = 0.085).

In order for small ncRNAs to function in gene expression control they must associate with proteins of the Argonaute family^49^. We screened immunoprecipitation data from all *C. elegans* AGOs^18^ to test whether tRNA 3’ halves could be found associated with specific AGOs. Both ERGO1 and WAGO10 showed a high enrichment in tRNA 3’ halves relative to control IPs (Fig.4.F). Specifically, a significant decrease in the number of Glutamate tRNAs fragments was detected between wild worms (N2) and the WAGO10 mutant which may contribute to the overall modest reduction in total tRNA fragments observed in the WAGO10 mutant. (Sup. Fig. 3. A & B).

We wondered whether tRNA 3’ halves induced by cisplatin treatment showed the characteristics of epimutations, in other words showing heritability over a short number of generations similarly to 22G-RNAs. We first compared the survival of 22G-RNAs and tRNAs, which demonstrated that tRNAs and 22G-RNAs were inherited for similar duration (Fig.4.G). We later observed a significant increase in the rate of tRNAs new epimutations in cisplatin high dose (Kruskal-Wallis rank sum test followed by pairwise Wilcoxon test with Bonferroni correction: HD vs. C: p-value = 0.003649917, HD vs. LD: p-value = 0.020226741) (Fig.4.H), suggesting that the overall increase in reads from tRNA halves is due to increased fluctuations in their expression rather than a consistent change. We then created a table with the number and maximal duration of epimutations by tRNAs type (Sup. Table 7). Several tRNAs were epimutated only in presence of cisplatin (TrpCCA, SerCGA, LeuAAG, GlyGCC, GlnCTG, AspGTC and AlaAGC). The two tRNAs with the highest number of epimutations were both found in high dose cisplatin and were GlyGCC and GlnTTG (Sup. Fig. 4. A & B).

However, the epimutations that occurred did not last longer than those occurring in control conditions (Log-rank test, p-value= 0.081) (Sup. Fig. 4. C). Thus, cisplatin exposure led to increased fluctuations in expression of tRNA halves which were not transmitted between generations.

### 5. Fluctuations in tRNA 3’ halves are associated with changes in expression of potential target genes

Having established that tRNA 3’ halves show increased fluctuations in their levels under conditions of cisplatin exposure, we wondered whether this might have any consequences for gene expression. We searched for potential targets of tRNAs. Whilst there were no genes with any regions demonstrating perfect complementarity to tRNA 3’ halves, we identified 38 genes with up to 2 mismatches. In all conditions, changes in expression of these genes were strongly associated with simultaneous changes in expression of tRNAs (Table 2, Test 1). Although the association was reduced in high dose cisplatin, due to the higher number of tRNAs fragments showing changes, the total number of genes showing changes in both tRNA fragments and gene expression was 1.4-fold higher in high dose cisplatin than in the control condition. In the control condition, association between gene expression change and tRNA 3’ halves epimutation was mostly discordant such that increased tRNA levels was associated with reduced gene expression (Fig.5.A). However, tRNA fragment increases were equally likely to be associated with up and down regulation of the genes under both high and low dose cisplatin exposure.

**Figure 5:**
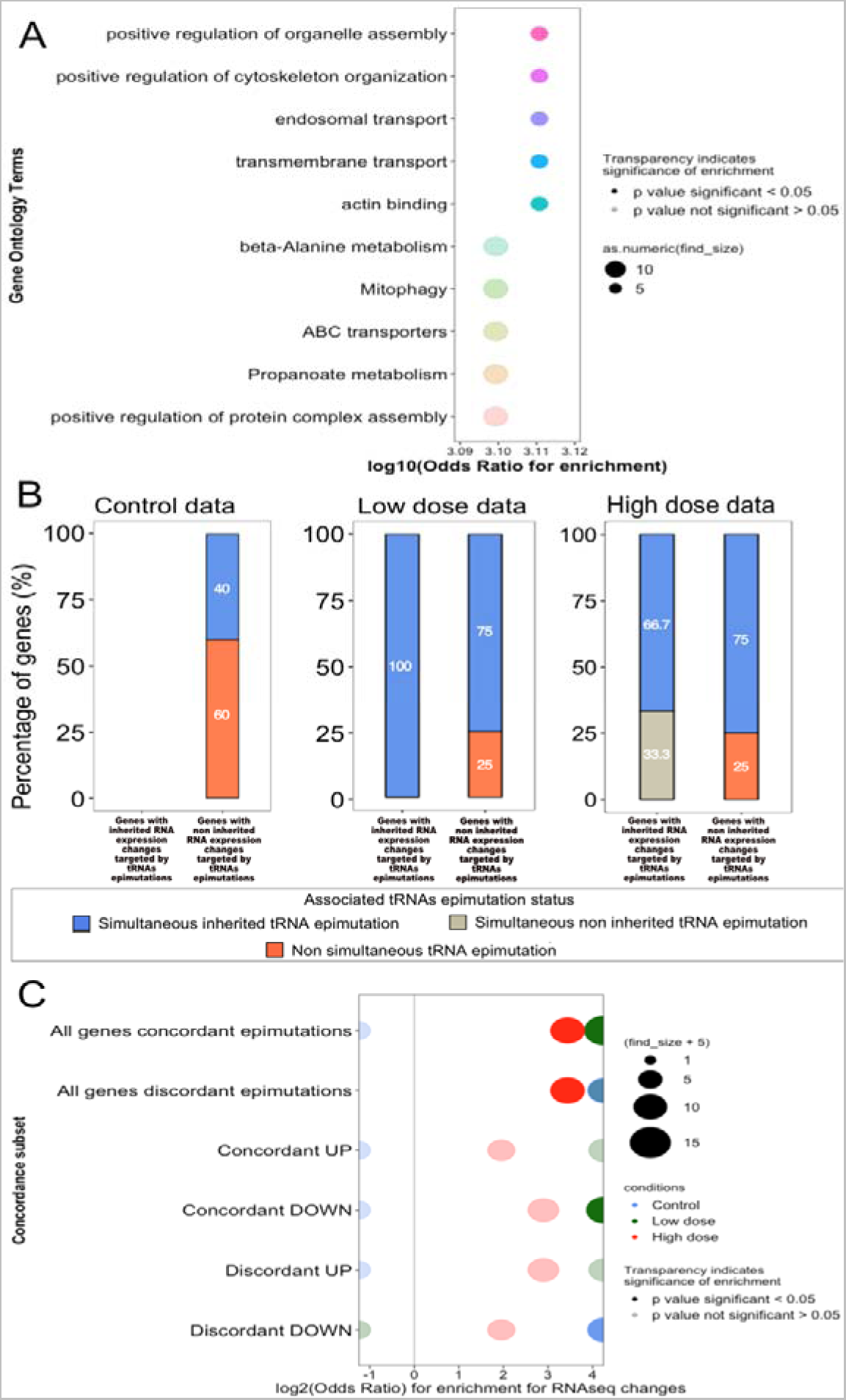
Correlation gene expression changes and tRNAs epimutations. A. Bubble plot showing ontology term enrichment of genes associated with tRNAs. Enrichment calculated with _χ_-squared test; top 10 results shown. X-axis showslog10(_χ_) for enrichment. Y-axis shows ontology terms. B. Stacked barplots representing the proportion of genes with inherited (left-bar) of non-inherited (right-bar) RNA expression changes targeted by tRNAs epimutations in control condition (left panel), cisplatin low dose condition (middle panel) and cisplatin high dose condition (right panel). In blue the percentage of genes with simultaneous inherited tRNAs epimutations, in grey the percentage of genes with simultaneous non-inherited tRNAs epimutations and in orange the percentage of genes with non-simultaneous tRNAs epimutations. C. Bubble plot of concordance (simultaneous and direction matched) or discordance (simultaneous but direction unmatched) of gene expression changes and tRNAs epimutations in control condition (blue dots), cisplatin low dose (green dots) or cisplatin *high dose* (red dots). Y-axis shows associations testes. X-axis shows log2(Odds ratio) of gene expression changes to be associated with tRNAs epimutation within each association. Odds ratios and p-values were calculated using Fisher’s Exact Test.

**Table 2:** Stepwise analysis of association of gene expression changes and tRNAs epimutations in the different exposure conditions. Cells in yellow indicate significantly increased odds of association, cells in grey indicate no significant association. Odds ratios and p-values were calculated with Fisher’s Exact Test.

| Condition | Test | Association tested | Background | Odds Ratio | p-value |
| --- | --- | --- | --- | --- | --- |
| Control | 1 | Gene exp. change & tRNA epimutations | All genes | Infinite | 0.0009587187 |
| Control | 2 | Inherited gene exp. change & tRNA epimutations | All genes with RNA-seq changes | 0.343171 | 0.03087491 |
| Control | 3 | Inherited gene exp. change & simultaneous tRNA epimutations | All genes with RNA-seq changes | 0 | 1 |
| Control | 4 | Inherited gene exp. change & simultaneous tRNA epimutations | All genes with RNA-seq changes and tRNA epimutations | 0 | 1 |
| Low dose | 1 | Gene exp. change & tRNA epimutations | All genes | Infinite | 8.862106e-05 |
| Low dose | 2 | Inherited gene exp. change & tRNA epimutations | All genes with RNA-seq changes | 0.6494231 | 1.000000e+00 |
| Low dose | 3 | Inherited gene exp. change & simultaneous tRNA epimutations | All genes with RNA-seq changes | 0.2345433 | 7.300642e-03 |
| Low dose | 4 | Inherited gene exp. change & simultaneous tRNA epimutations | All genes with RNA-seq changes and tRNA epimutations | Infinite | 1.000000e+00 |
| High dose | 1 | Gene exp. change & tRNA epimutations | All genes | 19.777393 | 0.0002813075 |
| High dose | 2 | Inherited gene exp. change & tRNA epimutations | All genes with RNA-seq changes | 1 | 1 |
| High dose | 3 | Inherited gene exp. change & simultaneous tRNA epimutations | All genes with RNA-seq changes | 1.605992 | 0.6979 |
| High dose | 4 | Inherited gene exp. change & simultaneous tRNA epimutations | All genes with RNA-seq changes and tRNA epimutations | Infinite | 1 |

We then looked at the percentage of genes with inherited expression change that were also targeted by changes in tRNA 3’ halves levels in each condition (Fig. 5.C). As for all genes with both gene expression and tRNA change, the percentage of genes with inherited gene expression change and simultaneous tRNA changes was lower in high dose than in the two other conditions (Table 2, test 2-4).

Interestingly, on average, 45.8% of changes in tRNA level were associated with a simultaneous gene expression change, compared to 8% for 22G-RNAs (Table 3). Moreover, some of the tRNA changes were associated with changes in gene expression that persisted for multiple generations (Fig.6).

**Figure 6:**
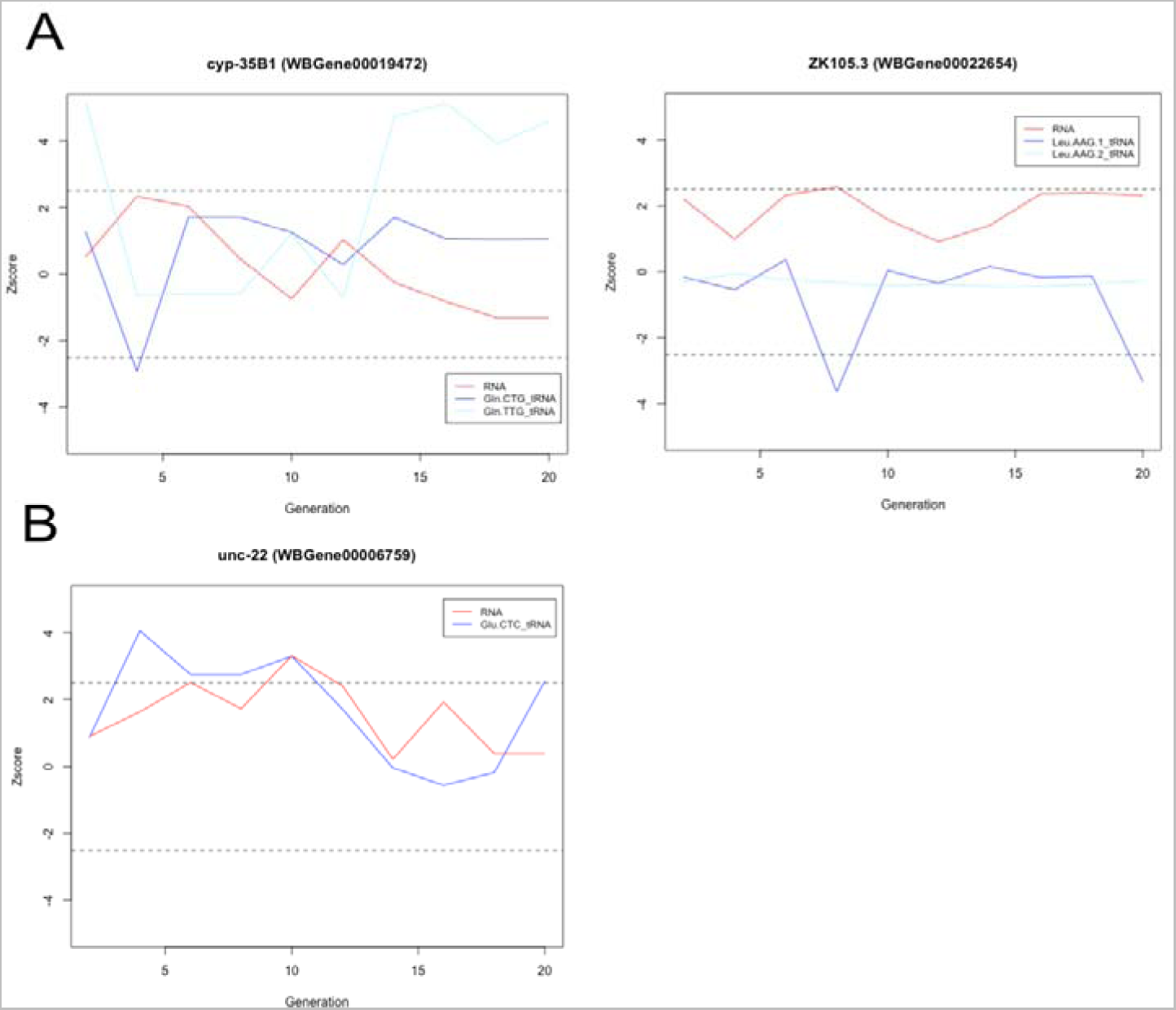
A. Two examples of genes showing an anticorrelation between gene expression change and tRNA 3’ halves epimutation. X-axe represent the generation, Y-axe shows z-score. Red line represents gene expression, blues lines represent tRNA 3’ halves. B. One examples of a gene showing a correlation between gene expression change and tRNA 3’ halves epimutation. X-axe represent the generation, Y-axe shows z-score. Red line represents gene expression, blue line represents tRNA 3’ halves.

**Table 3:** Comparison of odds of association between 22G-RNAs or tRNAs epimutations and gene expression changes. Cells in yellow indicate significantly increased odds of association for tRNAs compared to 22G-RNAs. Odds ratios and p-values were calculated with Fisher’s Exact Test.

| Condition | Association tested | X =22G-RNAs | X =tRNAs | Odds ratio | p-value |
| --- | --- | --- | --- | --- | --- |
| Control | genes with exp. change + simultaneous X epimutation | 75 | 2 | 0.6582255 | 0.6422611 |
| Control | genes with exp. change only | 912 | 16 |  |  |
| Low dose | genes with exp. change + simultaneous X epimutation | 73 | 4 | 0.15656273 | 0.009301 |
| Low dose | genes with exp. change only | 936 | 8 |  |  |
| High dose | genes with exp. change + simultaneous X epimutation | 65 | 6 | 0.1024083 | 0.000296 |
| High dose | genes with exp. change only | 851 | 8 |  |  |

One possible mechanism whereby tRNA 3’ half changes could result in gene expression changes could be due to induction of 22G-RNAs downstream of recognition of target genes by tRNA-Argonaute complexes. However, most genes targeted by tRNA fragments did not have 22G-RNA mapping to them, thus the mechanism whereby tRNA 3’ halves result in gene expression changes in *C. elegans* remains to be determined.

## Discussion

As previously discussed in the literature^39–41^, our results confirmed that cisplatin exposure led to genotoxic stress with an increase in the rate of random mutations. We wanted to investigate if cisplatin might also affect the rate, spectrum, or duration of epimutations. Our results showed that epimutations were not strongly affected by cisplatin exposure, as identified changes were similar in both control and cisplatin conditions. However, cisplatin tends to increase the expression of reads mapping to 3’ tRNA halves. We discuss these results in terms of their significance for the contribution of epimutations to evolutionary processes and potential new insights into the biology of small non-coding RNAs in *C. elegans*.

Previous work has explored the rate, spectrum and stability of epimutations in plants^13–15, 50, 51^ and *C.elegans*^9, 11^ in the absence of selection, demonstrating that epimutations arise much more frequently that genetic changes involving DNA sequence alterations, but have a limited duration. This suggests that epimutations are unlikely to contribute significantly to genetic drift over long periods of time^14, 52^. However, this still leaves open the possibility that epimutations may contribute to evolution under conditions of strong selection^53^. The selection pressure would have to be strong enough to outweigh the limited half-life of any beneficial epimutation due to their limited stability. A key unknown in this discussion, nonetheless, is whether the rate of occurrence, the stability, or the type of epimutations that occur is independent of the nature of the selective pressure itself. Our results pertain to this assumption, using genotoxic stress as an example of a selective pressure. We measured rate, stability, and spectrum of epimutations in both gene expression and small non-coding RNAs, the pre-eminent molecular mechanism that gives rise to transgenerational epigenetic inheritance in *C. elegans* in the absence of selection, whilst applying genotoxic stress through cisplatin exposure. Crucially, under these conditions we saw marginal changes in the global parameters of gene expression and small non-coding RNAs epimutations. This is an important result for evaluating the potential for such kind of epimutations to stimulate evolution as it shows that, at least for cisplatin exposure, epimutations are neither increased in overall rate nor directed to specific regions of the genome that might have adaptative consequences. The idea that epimutations might respond to environmental changes and thus provide particular significance in adaptation has been frequently proposed^54–57^, and some authors have argued that this might operate in plants^58^. Our results argue against this proposal, at least in the case of small non-coding RNAs-mediated epimutations in cisplatin exposure in *C. elegans*. Systematic analyses of epimutations in the absence of selection under different stresses would be required to comprehensively evaluate this idea.

Whilst we did not observe changes in the parameters of epimutations in response to cisplatin, we did observe changes in the levels of tRNA-derived small non-coding RNAs. Work across a range of model systems have identified tRNA-derived small RNAs as an abundant component of the cellular small non-coding RNA fraction^59–64^. Most commonly these are either 5’ or 3’ halves and arise through cleavage of the tRNA by several enzymes, most prominently RNAses^65–70^. Even if this appears to happen constitutively in mammalian cultured cells^71^, tRNA fragments are further induced by cellular stress, including oxidative stress^72^ and infection^73^.

Despite their widespread conservation^74^, tRNA fragments have not been extensively characterised in *C. elegans*. Our results demonstrate a clear increase in the levels of tRNA 3’ halves associated with exposure to cisplatin. The fact that RNA 3’ halves become much more abundant upon genotoxic stress may explain why tRNAs were not previously noted as abundant features of *C. elegans* small RNA populations. Interestingly, one of the few publications describing tRNA fragments in *C. elegans* demonstrated an increased abundance of 3’ tRNA fragments in aged worms^75^, which support the idea that they may be rare in unstressed worms but increase dramatically under certain stressful conditions. Moreover, this is consistent with their induction by cellular stress in mammalian cells.

What might be the function of the increased levels of tRNA halves in response to cisplatin? Stress-induced cleavage of tRNAs in mammalian cells is associated with translational repression^72, 76^ and this may be a function of tRNA cleavage in cisplatin-exposed *C. elegans*. Indeed, the fact that the changes in tRNA fragments levels are highly variable across different cisplatin-exposed lines even within the same lineage suggest a response to cellular stress, potentially shutting down translation in cells that experience particularly high levels of damage. It may also be the case that the tRNA halves themselves have direct function in gene expression control^77^. Our data suggest that there may be a role for the argonaut protein WAGO10 in binding and stabilising tRNA 3’ fragments. tRNA fragments might therefore recruit WAGO10 to mRNAs based on imperfect sense-antisense base pairing and indeed, we determined a significant tendency for the levels of tRNAs 3’ halves to be associated with changes in expression of potential targets. Association of tRNA fragments with Argonauts proteins has been documented in mammalian cells^74, 78–80^ and plants^59, 81^; however, whether this results in gene expression changes is still debated. *C. elegans* may thus be a good model for future investigations of the mechanistic basis of tRNA fragments in gene expression control.

Our discovery that tRNA fragments appear to be induced by cisplatin exposure in *C. elegans* is interesting in the light of several recent studies showing that tRNA fragments can be transmitted intergenerationally through sperm in rodents^4, 48, 82–85^. Although alterations in tRNA fragments levels that we observed upon cisplatin treatment were mostly transient and not inherited transgenerationally, we did observe a small number of examples of epimutations involving tRNA fragments and their targets. Mechanistically, transgenerational epigenetic inheritance of small non-coding RNAs in *C. elegans* relies on RNA-dependent RNA polymerase (RdRP) to amplify the response each generation^86, 87^. It is unlikely that RdRPs could copy tRNA fragments, so tRNA fragments are likely to only survive for a very small number of generations on average, which is consistent with our observations. In addition, we did not see evidence of induction of 22G-RNAs mapping to tRNA targets. It is not therefore our current view that tRNA fragments induction acts as a significant source of epimutations that is revealed upon cisplatin treatment. Nevertheless, future work using stress to induce tRNA derived fragments in *C. elegans* could reveal new mechanisms whereby this could lead to transgenerational epigenetic changes in gene expression that persist more than one or two generations.

## Data availability

The sncRNAs data have been deposited with links to BioProject accession number PRJNA991138 in the NCBI BioProject database (https://www.ncbi.nlm.nih.gov/bioproject/)

All code for the R analysis including figure plotting commands is on the GitHub page.

## Declaration of Competing Interest

The authors declare that they have no known competing financial interests or personal relationships that could have appeared to influence the work reported in this paper.

## Acknowledgements

This project was supported by the EPA Cephalosporin fund granted to Peter Sarkies. We thank members of the Epigenetic and Evolution group for fruitful discussions.

## Author Contributions

Conceptualization: Rachel Wilson, Peter Sarkies.

Formal analysis: Manon Fallet, Peter Sarkies.

Investigation: Manon Fallet, Peter Sarkies.

Methodology: Manon Fallet, Rachel Wilson, Peter Sarkies.

Supervision: Peter Sarkies.

Writing – original draft: Manon Fallet, Rachel Wilson, Peter Sarkies

Writing – review & editing: Manon Fallet, Rachel Wilson, Peter Sarkies

## Supplementary Files Legends

**Sup.Fig.1: Additional pieces of evidence supporting the genotoxic effect of cisplatin on worms**. A. Boxplot of the number of worms in control or in cisplatin high dose condition. No significant difference was observed between the two conditions (T-test, pval=0.05714). B. Bubble plot showing ontology term enrichment of genes with gene expression changes in high dose cisplatin compared to genes without expression change. Enrichment calculated using Fisher’s Exact Test with Bonferroni correction, top 10 results are shown, X-axis shows log10(Odds) or enrichment. Y-axis shows ontology terms. P-value cut-off for significance is 0.05.

**Sup. Fig. 2: Association between 22G-RNAs epimutations and gene expression epimutations.** A. Stacked barplots showing the percentage of genes with inherited (left-bar) of non-inherited (right-bar) RNA expression changes targeted by 22G-RNAs epimutations in control condition (left panel), cisplatin low dose condition (middle panel) and cisplatin high dose condition (right panel). In blue the percentage of genes with simultaneous inherited 22G-RNAs epimutations, in grey the percentage of genes with simultaneous non-inherited 22G-RNAs epimutations and in orange the percentage of genes with non-simultaneous 22G-RNAs epimutations.

**Sup. Fig. 3: Association between tRNAs and specific AGOs proteins.** A. Plot of total tRNAs count (in reads per million) in wild type C. elegans (N2; black diamonds) and mutants lacking specific AGOs proteins (from left to right: ergo1 (blue diamonds), rde1 (yellow diamonds) and wago10 (red diamonds). A very small decrease in wago10 was observed. B. Plot representing specifically Glutamate tRNA (GluCTC (diamonds) and GluTCC (dots) fragments (in reads per million) in wild type C. elegans (N2; black) and in mutants lacking wago10 (red). A significant difference in Glutamate tRNA fragments was observed between wild type worms and mutants lacking wago10 (Wilcox-test, adjusted p-value = 0.048).

**Sup. Fig. 4: Detailed characterisation of tRNAs epimutations.** A. Barplot of total number of epimutations for each tRNAs type and in the different conditions: control (blue), LD (green) and HD (red). B. Barplot of the duration of tRNAs epimutations for each kind and according to the exposure condition: control (blue), LD (green) and HD (red). C. Forest plot of Cox Proportional Hazards Model representing the odd of difference in the tRNAs 3’ halves epimutations between the conditions. The x-axis shows the chances of an epimutation to disappear in the cisplatin conditions in comparison to control (reference). The p-values were calculated using log rank test.

**Sup. Table. 1. Reference table for sncRNAs type identification in the tinyRNA pipeline.**

**Sup. Table. 2. Reads count number for sncRNAs mapping to tRNAs.** Readcount for each small RNA mapping to a tRNA was normalized using the size factors from DESeq calculated as part of the tinyRNA pipeline. Counts for individual tRNA types were obtained by summing normalized reads mapping to the 3’ half of each tRNA type.

**Sup. Table. 3. tRNAs 3’ halves potential target genes.** To identify potential target genes, small RNAs corresponding to tRNA 3’ halves were extracted from all the lines and combined into a single file, along with the annotation indicating which tRNA the sequence was derived from. This was aligned to the ce11 genome allowing up to 3 mismatches. Protein-coding genes overlapping with mismatched tRNA fragments were then obtained using bedtools intersect, to identify genes that were potentially targeted by tRNAs.

**Sup. Table. 4. Gene ontology enrichment result for high dose cisplatin.** 279 significantly enriched functions with genes expression changes in high dose cisplatin condition compared to control were identified using DESeq2 and EnrichR.

**Sup. Table. 5. Gene ontology analysis for gene expression epimutations.** Gene ontology analysis results for gene expression changes across the mutation accumulation lines.

**Sup. Table. 6. Stepwise analysis of association of gene expression changes and 22G-RNAs epimutations in the different exposure conditions.** Cells in yellow indicate significantly increased odds of association, cells in grey indicate no significant association. Odds ratios and p-values were calculated with Fisher’s Exact Test.

**Sup. Table. 7. Number and maximal duration of epimutations by tRNAs type.**

## Notes

### Competing Interest Statement

The authors have declared no competing interest.

