## Supplemental Figure1 for "Chronic cisplatin exposure does not affect epimutations in *C. elegans* but induces fluctuations in tRNA-derived small non-coding RNAs"

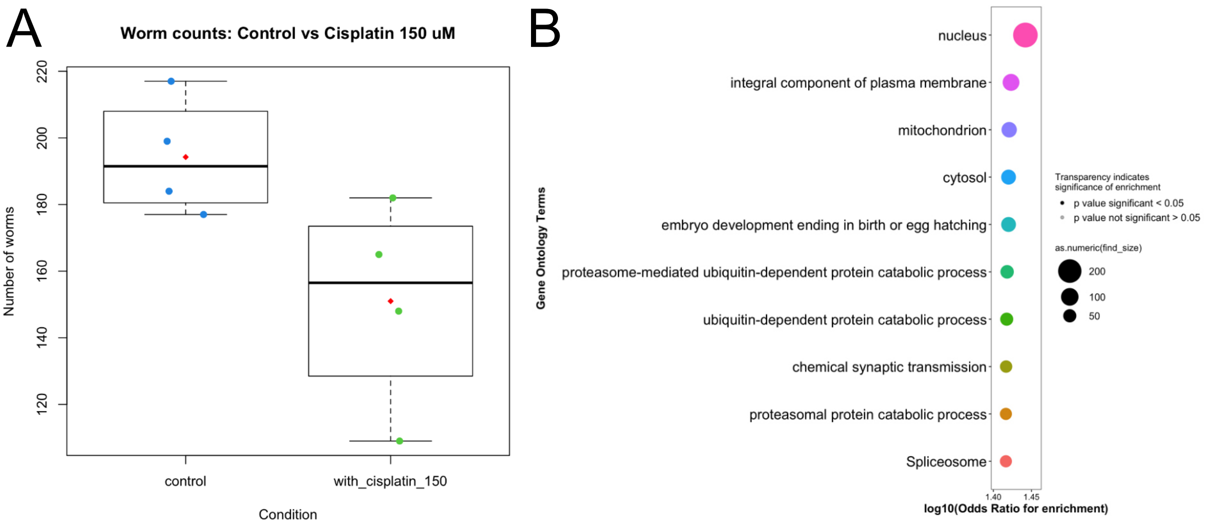


**Sup.Fig.1: Additional evidences supporting the genotoxic effect of cisplatin on worms.** A. Boxplot of the number of worms in control or in cisplatin high dose condition. No significant difference was observed between the two conditions (T-test, pval=0.05714). B. Bubble plot showing ontology term enrichment of genes with genes expression changes in high dose cisplatin compared to genes without expression change. Enrichment calculated using Fisher's Exact Test with Bonferroni correction, top 10 results shown, X-axis shows log10(Odds) or enrichment. Y-axis shows ontology terms. P-value cut off for significance is 0.05.
