## Supplemental Figure 2 for "Chronic cisplatin exposure does not affect epimutations in *C. elegans* but induces fluctuations in tRNA-derived small non-coding RNAs"

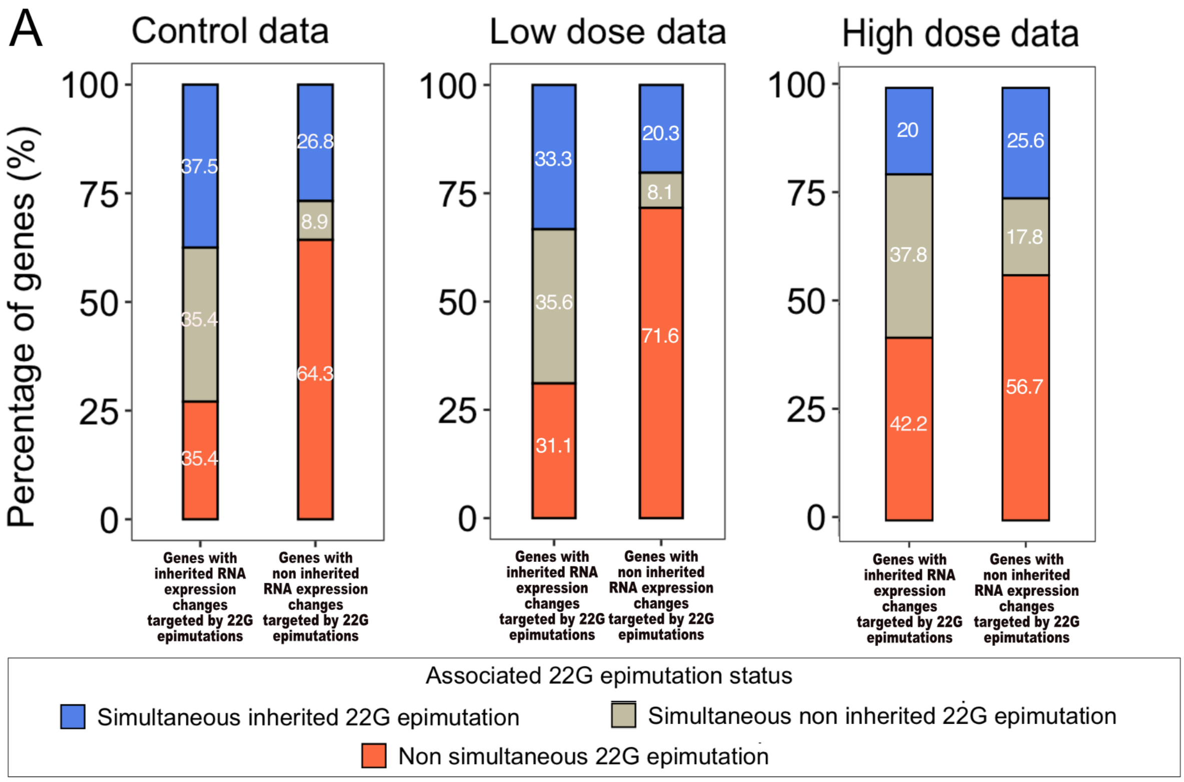


**Sup. Fig. 2: Association between 22G-RNAs epimutations and gene expression epimutations.** A. Stacked barplots showing the percentage of genes with inherited (left-bar) of non-inherited (right-bar) RNA expression changes targeted by 22G-RNAs epimutations in control condition (left panel), cisplatin low dose condition (middle panel) and cisplatin high dose condition (right panel). In blue the percentage of genes with simultaneous inherited 22G-RNAs epimutations, in grey the percentage of genes with simultaneous non-inherited 22G-RNAs epimutations and in orange the percentage of genes with non-simultaneous 22G-RNAs epimutations.
