## Supplemental Figure 3 for "Chronic cisplatin exposure does not affect epimutations in *C. elegans* but induces fluctuations in tRNA-derived small non-coding RNAs"

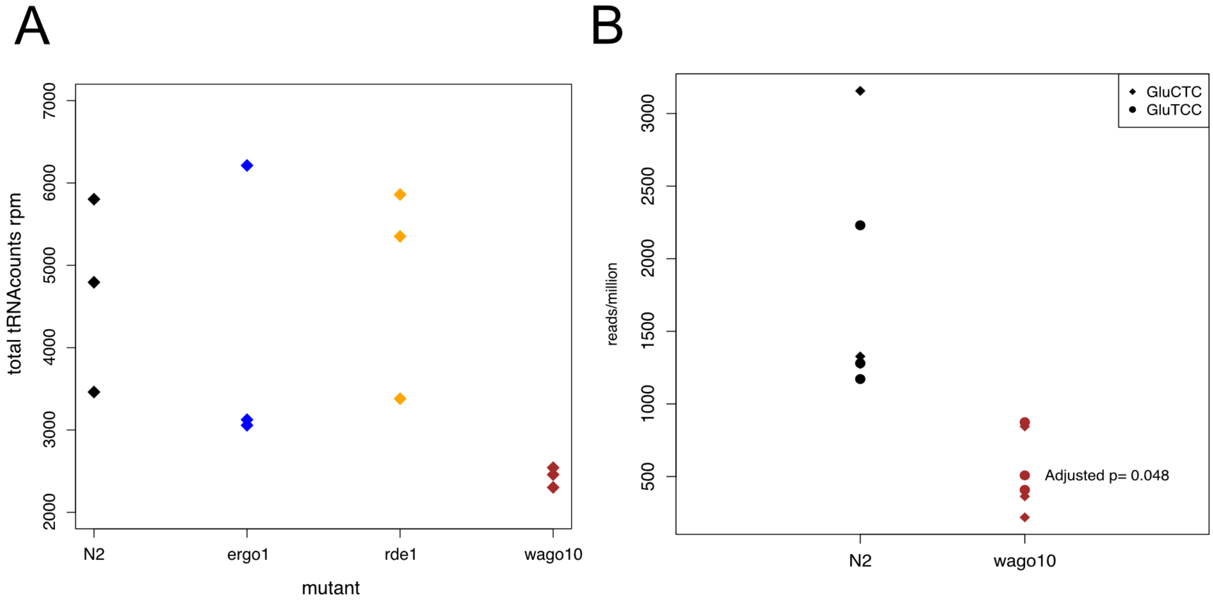


**Sup. Fig. 3: Association between tRNAs and specific AGOs proteins.** A. Plot of total tRNAs count (in reads per million) in wild type *C. elegans* (N2; black diamonds) and mutants lacking specific AGOs proteins (from left to right: ergo1 (blue diamonds), rde1 (yellow diamonds) and wago10 (red diamonds). A very small decrease in wago10 was observed. B. Plot representing specifically Glutamate tRNA (GluCTC (diamonds) and GluTCC (dots) fragments (in reads per million) in wild type *C. elegans* (N2; black) and in mutants lacking wago10 (red). A significant difference in Glutamate tRNA fragments was observed between wild type worms and mutants lacking wago10 (Wilcox-test, adjusted p-value = 0.048).
