## Supplemental Figure 4 for "Chronic cisplatin exposure does not affect epimutations in *C. elegans* but induces fluctuations in tRNA-derived small non-coding RNAs"

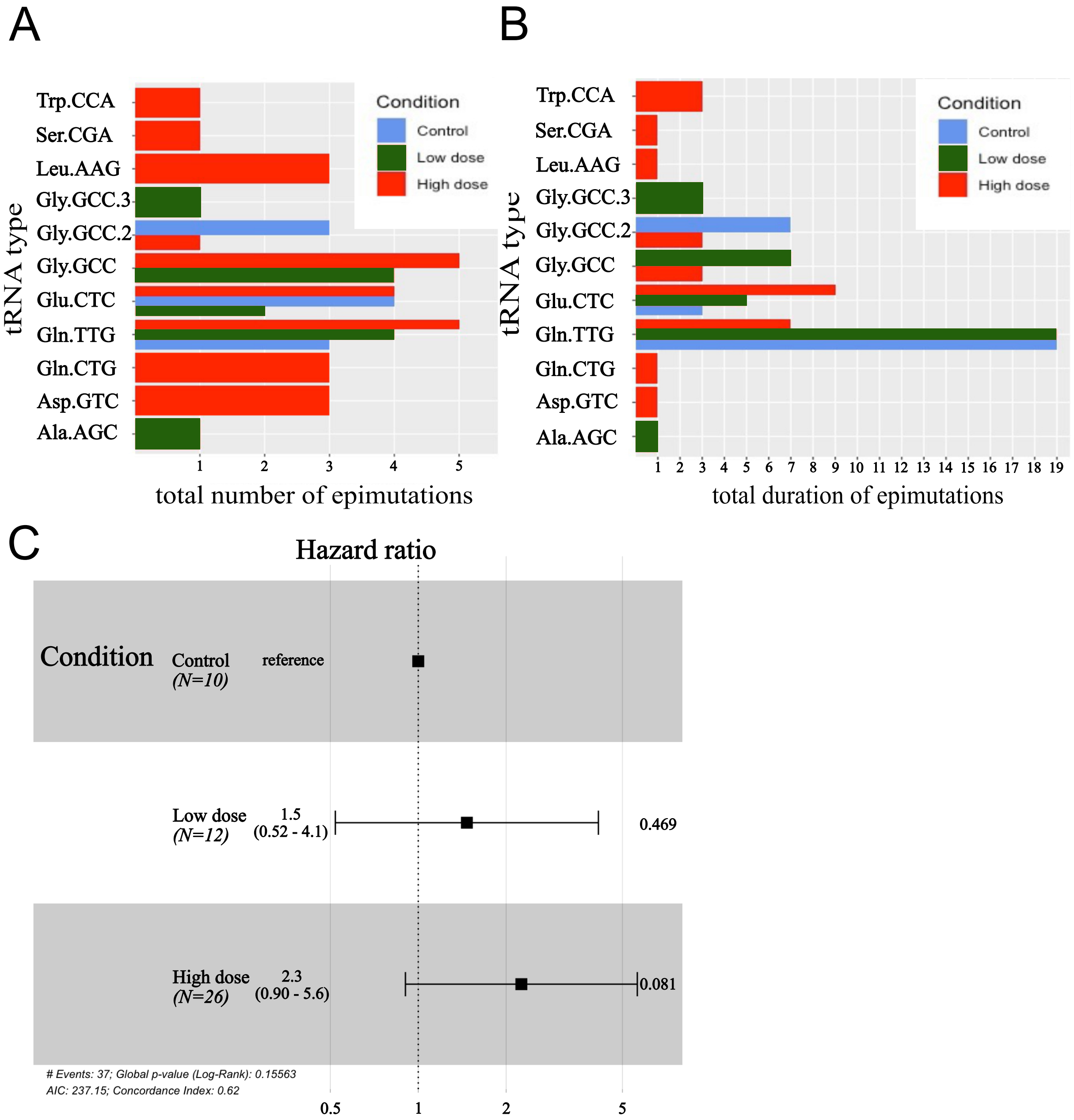


**Sup. Fig. 4: Detailed characterisation of tRNAs epimutations.** A. Barplot of total number of epimutations for each tRNAs type and in the different conditions: control (blue), LD (green) and HD (red). B. Barplot of the duration of tRNAs epimutations for each kind and according to the exposure condition: control (blue), LD (green) and HD (red). C. Forest plot of Cox Proportional Hazards Model representing the odd of difference in the tRNAs 3' halves epimutations between the conditions. The x-axis show the chances of an epimutation to disappear in the cisplatin conditions in comparison to control (reference). The p-values were calculated using log rank test.
