## Supplemental Table 3 for "Chronic cisplatin exposure does not affect epimutations in *C. elegans* but induces fluctuations in tRNA-derived small non-coding RNAs"

| Name | seqID | AA | codon | mismatch |
|---|---|---|---|---|
| Y67H2A.6b.1 | IV:620502-620521(+) | Val | CAC | 1 |
| F23A7.8.1 | I:6135829-6135848(+) | Tyr | GTA | 1 |
| M03C11.6.1 | I:9267150-9267169(+) | Arg | TCG | 1 |
| ZK105.3.1 | I:6051154-6051173(-) | Leu | AAG | 1 |
| ZK105.3.1 | II:6721829-6721848(+) | Leu | TAA | 1 |
| T06D8.1g.1 | I:2272406-2272427(-) | Gly | GCC | 2 |
| F09F7.4a.1 | I:12142946-12142965(+) | Met | CAT | 2 |
| K07C6.4.1 | II:7756967-7756987(-) | Gln | TTG | 2 |
| F54C9.5.1 | III:4362523-4362543(+) | Leu | CAA | 2 |
| K07C6.4.1 | IV:5334745-5334764(+) | Gln | CTG | 2 |
| F23A7.4.1 | I:6135826-6135848(+) | Tyr | GTA | 2 |
| M03C11.6.1 | I:9267151-9267171(+) | Arg | TCG | 2 |
| Y7A5A.1.1 | I:11803315-11803334(-) | Ser | CGA | 2 |
| Y7A5A.1.1 | II:1520097-1520116(+) | Ser | AGA | 2 |
| Y49F6C.3.1 | MtDNA:1666-1685(+) | NA | NA | 2 |
| ZK617.1b.1 | III:7642179-7642198(-) | Glu | CTC | 2 |
| B0432.11.1 | IV:144018-144037(+) | Arg | CCT | 2 |
| W01A8.5.1 | IV:620518-620537(+) | Val | CAC | 2 |
| ZK652.2.1 | IV:16577337-16577356(+) | Ser | TGA | 2 |
| C49F8.3.1 | I:6136638-6136657(+) | Lys | CTT | 2 |
| F14F4.3.1 | I:12065878-12065897(+) | Val | AAC | 2 |
| C56G3.1a.2 | I:9320004-9320023(-) | Gly | TCC | 2 |
| Y67H2A.6b.1 | II:5577232-5577251(-) | Val | TAC | 2 |
| F55G1.4.1 | II:10302204-10302225(-) | Pro | TGG | 2 |
| C01B4.10.1 | V:7730670-7730689(-) | Thr | CGT | 2 |
| F52C9.1.1 | I:6781850-6781870(+) | Cys | GCA | 2 |
| T09D3.1.1 | I:5854837-5854856(-) | Pro | AGG | 2 |
| Y43F8B.17.1 | I:8953210-8953229(+) | Lys | TTT | 2 |
| C05D11.2.1 | I:5843094-5843113(-) | Arg | ACG | 2 |
| F23A7.8.1 | I:10945820-10945840(+) | Phe | GAA | 2 |
| F55A11.4a.1 | II:3439218-3439237(-) | Thr | AGT | 2 |
| T07F10.1b.1 | II:11334023-11334042(-) | Ala | TGC | 2 |
| Y94A7B.5.1 | IV:13870269-13870291(+) | His | GTG | 2 |
| C14C10.3c.1 | II:5772976-5772995(-) | Leu | TAA | 2 |
| Y73F8A.2.1 | X:12890301-12890320(-) | Ala | CGC | 2 |
| F02D10.3.1 | I:1447168-1447188(+) | Arg | TCT | 2 |
| F41E7.5.1 | I:13310354-13310373(-) | Thr | TGT | 2 |
| Y110A7A.7.1 | IV:658245-658265(+) | SeC(e) | TCA | 2 |
| T06D8.1d.1 | I:2272406-2272427(-) | Gly | GCC | 3 |
| K09H11.3.1 | I:6051217-6051236(-) | Leu | AAG | 3 |
| F23A7.4.1 | I:6135826-6135850(+) | Tyr | GTA | 3 |
| C26C6.5b.1 | I:6163524-6163544(-) | Ile | AAT | 3 |
| T22F3.10.1 | I:8530149-8530168(-) | Glu | CTC | 3 |
| T06H11.1a.1 | I:9267141-9267161(+) | Arg | TCG | 3 |
| C28D4.9.1 | I:9320005-9320025(-) | Gly | TCC | 3 |
| ZK970.1b.1 | I:9883726-9883745(+) | Glu | TTC | 3 |
| C05D11.5.1 | I:10807225-10807244(+) | Asp | GTC | 3 |
| Y71F9AL.16.1 | I:11803326-11803345(-) | Ser | CGA | 3 |
| M142.6c.1 | I:12065907-12065926(+) | Val | AAC | 3 |
| F09F7.4b.1 | I:12142946-12142965(+) | Met | CAT | 3 |
| C50F4.13.1 | I:12601814-12601834(+) | Ala | AGC | 3 |
| Y71F9AL.16.1 | II:1520086-1520105(+) | Ser | AGA | 3 |
| Y45G12B.1.1 | II:5577218-5577238(-) | Val | TAC | 3 |

| | | | | | |
|---|---|---|---|---|---|
| F46H5.4.1 | II:6721765-6721784(+) | Leu | TAA | 3 | |
| C50F4.13.1 | II:7003313-7003333(+) | Thr | AGT | 3 | |
| C17H1.7.1 | II:7756969-7756989(-) | Gln | TTG | 3 | |
| F17B5.2.1 | II:10302205-10302226(-) | Pro | TGG | 3 | |
| Y45G12B.1.1 | IV:620515-620535(+) | Val | CAC | 3 | |
| F52B11.3.1 | IV:5334742-5334764(+) | Gln | CTG | 3 | |
| T26E4.4.1 | MtDNA:91-111(+) NA | NA | 3 | | |
| F47C12.2.1 | X:6371375-6371396(+) | Trp | CCA | 3 | |
| C26C6.5b.1 | X:9059374-9059394(-) | Undet | ??? | 3 | |
| Y38C1AA.9.1 | I:754740-754759(+) | Leu | CAG | 3 | |
| Y43F8B.17.1 | I:8953209-8953229(+) | Lys | TTT | 3 | |
| C41C4.4a.1 | I:9051306-9051325(-) | Asn | GTT | 3 | |
| Y51A2D.17.1 | I:14162845-14162866(+) | Ser | GCT | 3 | |
| Y67D2.6.1 | II:3521508-3521527(-) | His | GTG | 3 | |
| T27E9.7.1 | III:8650419-8650438(-) | Pro | CGG | 3 | |
| T01G5.5.1 | V:12780447-12780469(-) | Ser | TGA | 3 | |
| F55G1.4.1 | X:3970101-3970123(+) | Pro | AGG | 3 | |
| Y95B8A.5a.1 | X:6220086-6220105(-) | Arg | ACG | 3 | |
| K10D6.2d.1 | I:1447173-1447193(+) | Arg | TCT | 3 | |
| F31C3.3.1 | I:6136635-6136656(+) | Lys | CTT | 3 | |
| K03A11.6.1 | I:13310357-13310376(-) | Thr | TGT | 3 | |
| T19D12.5.1 | II:11334031-11334050(-) | Ala | TGC | 3 | |
| F33H12.5.1 | III:11792815-11792834(+) | | Thr | CGT | 3 |
| T22H9.3.1 | IV:144022-144043(+) | Arg | CCT | 3 | |
| T19D12.5.1 | V:9503595-9503614(+) | Ala | CGC | 3 | |
| F21F8.3.1 | I:6781838-6781857(+) | Cys | GCA | 3 | |
| Y69H2.3b.1 | V:6917527-6917546(+) | Ile | TAT | 3 | |
| F23A7.8.1 | III:7978523-7978545(-) | Phe | GAA | 3 | |
| W02A2.5.1 | IV:658181-658202(+) | SeC(e) | TCA | 3 | |
| Y105C5B.28c.1 | II:12728796-12728815(-) | Arg | CCG | 3 | |
| C14C10.3c.1 | II:6853203-6853222(+) | Leu | TAG | 3 | |
